# Data-independent immunopeptidomics discovery of low-abundant bacterial epitopes

**DOI:** 10.1101/2025.05.14.653952

**Authors:** Patrick Willems, An Staes, Vadim Demichev, Simon Devos, Francis Impens

**Author notes:** **Corresponding Author** Patrick Willems,; Francis Impens.

## Abstract

Mass spectrometry-based immunopeptidomics is a powerful approach to uncover peptides presented by human leukocyte antigen (HLA) molecules that can guide vaccine design and immunotherapies. While data-dependent acquisition (DDA) has been the standard for navigating through the complexity associated with non-enzymatic immunopeptide database searches, data-independent acquisition (DIA) is increasingly adopted in immunopeptidomics research. In this work, we compare diaPASEF to conventional ddaPASEF in terms of global immunopeptidome profiling and bacterial epitope discovery of the model intracellular pathogen *Listeria monocytogenes*. We show that DIA spectrum-centric workflows that search pseudo-MS/MS spectra complement DDA analysis by uncovering additional human and bacterial immunopeptides. Furthermore, we leveraged DIA-NN for generating and searching proteome-wide predicted HLA class I peptide spectral libraries, scoring approximately 150 million immunopeptide peptide precursors. This approach outperformed other spectrum-based methods in identification of MHC class I peptides and recovered low-abundant peptide precursors missed by other methods. Taken together, our results demonstrate how both DIA spectrum- and peptide-centric immunopeptidomics analysis are promising strategies to identify low-abundant immunopeptides.

## INTRODUCTION

Human leukocyte antigen (HLA) molecules present short peptide ligands or immunopeptides at the cell surface. HLA presentation of foreign bacterial epitopes of infected cells can be recognized by T-cell receptors of CD8+ T-cells to clear infected cells. As such, identification of these presented bacterial epitopes can elucidate antigens that can be used in vaccine formulation.^1^ Sensitive mass spectrometry methods enable to detect and quantify HLA-bound immunopeptides from purified fractions,^2^ and has been used by us and other groups to identify epitopes of various bacterial pathogens.^3-8^ We previously infected human cultured cells with *Listeria monocytogenes*, a foodborne model pathogen further referred to as *Listeria*, thereby revealing 64 bacterial epitopes and prioritizing multiple vaccine candidates.^4^ We recently improved immunopeptide detection by 76% in infected HeLa cell cultures using parallel accumulation–serial fragmentation (PASEF)^9^ on a timsTOF SCP instrument, levering four search engines and data-driven rescoring.^10^ Here, dual trapped ion mobility spectrometry (TIMS) allows to trap ions in a first analyzer, which are subsequently separated and released in terms of their ion mobility in the second analyzer, whilst synchronizing MS/MS precursor selection with TIMS separation. However, a more complete recording of precursor ions is achieved in diaPASEF acquisition, where predefined ion mobility and *m/z* windows are used to comprehensively and repeatedly sample the entire precursor space, with a potential of greater sensitivity and data completeness, as well as more accurate relative quantification.^11^

However, unlike conventional DDA, analysis of DIA data is tedious by complex MS/MS spectra of intermixed peptide precursors as peptides are co-fragmented. Furthermore, immunopeptide searches are associated with a large peptide search space, as opposed to regular tryptic digest analyses. To circumvent this, immunopeptidomics studies often restrict searches to predicted HLA peptide binders.^12,13^ Nevertheless, direct searches of DIA data against the full sequence database, performed in addition or instead of the analysis of predicted HLA complex binders, have the potential to maximize the number of HLA peptides discovered and quantified.

There are two main strategies to analyze DIA data and both have been used for immunopeptide identification. In peptide-centric approaches, spectral libraries are used that aim to identify peptide precursors in the library using provided retention time, fragment ion intensities and potentially other information such as ion mobility for each precursor that is provided in the library. These libraries can empirically rely on identifications of previous DDA runs, an approach used in several immunopeptidomics studies.^14-16^ Alternatively, proteome-wide spectral libraries can be generated *in silico* using deep learning models and are known to be highly performant for DIA analysis.^17,18^ Such predicted libraries are for example generated by the DIA-NN software, requiring solely a protein sequence database as input.^19,20^ An alternative strategy for DIA analysis are spectrum-centric approaches that deconvolute pseudo-MS/MS spectra from DIA data, which subsequently can be searched by conventional DDA search algorithms. This concept is originally used in DIA-Umpire^21^ and currently implemented for diaPASEF acquisition data analysis by the directDIA algorithm within Spectronaut^22^ and the diaTracer workflow recently introduced within FragPipe.^23^

We reasoned that complete and sensitive recording by diaPASEF could benefit the detection of low-abundant bacterial immunopeptides presented on infected cells. To this end, we evaluated how diaPASEF acquisition of *Listeria*-infected cell cultures facilitates the discovery of bacterial antigens, compared to ddaPASEF acquisition. Next to leveraging spectrum-centric analyses via diaTracer and directDIA, we demonstrate how proteome-wide *in silico* spectral libraries generated and searched by DIA-NN, using its ‘proteoforms’ confidence mode, recover low abundant immunopeptides missed by other approaches. Together, this showcases the promise of DIA analysis for immunopeptidomics-based discovery of bacterial antigens and other applications.

## EXPERIMENTAL PROCEDURES

### diaPASEF LC-MS/MS

Immunopeptides from *Listeria*-infected or uninfected HeLa cells were isolated as described.^4^ Leftover peptide mixtures peptides (9 µl) injected for ddaPASEF^10^ were supplemented with 6 µl loading solvent A (0.1% trifluoroacetic acid (TFA) in water/acetonitrile (ACN) (99.5:0.5, v/v)) of which 12 out of 15 µl was for LC-MS/MS analysis on an Ultimate 3000 RSLC nanoLC in-line connected to a timsTOF SCP mass spectrometer (Bruker). As in the ddaPASEF injection,^10^ this roughly approximates an equivalent of ∼30-50 million infected cells. Trapping was performed at 20 μl/min for 2 min in loading solvent A on a 5 mm trapping column (Thermo scientific, Pepmap, 300 μm internal diameter (I.D.), 5 μm beads). The sample was further separated on a reverse-phase column (Aurora elite 75µm x 150 mm 1.7um particles, IonOpticks) following elution from the trapping column by a linear gradient starting at 0.5% MS solvent B (0.1% FA in water/ACN 20:80 (v/v)) at a flow rate of 250 nl/min for 30 min reaching 37.5% MS solvent B, increasing MS solvent B to 55% MS solvent B after 38 min, finally increasing further to 70% MS solvent B after 40 min, followed by a wash for 5 min and re-equilibration with 99.5% MS solvent A (0.1% FA in water). The flow rate was decreased from 250 nl/min to 100 nl/min at 20 min and increased again to 250 nl/min at 40 min.

A control JY cell line immunopeptidomics sample was run in ddaPASEF and searched with MSFragger using the Nonspecific-HLA workflow with spectral library generation enabled. The created library was loaded into py_diAID^24^ to create the ion cloud. A diaPASEF schema was created with eight frames containing tree windows per PASEF scan (**Supplemental Table 1**). The schema was optimized through py_diAID and the schema covering most of the ion cloud was selected. Next manual adjustment was done to cover the ion cloud completely.

### Spectrum-centric searches

All data were searched against a custom concatenated database of the human (20,590 proteins) and *Listeria monocytogenes* EGD (2,847 proteins) UniProtKB reference proteomes. Previous ddaPASEF Comet, Sage and MSFragger search results^10^ were now supplemented with search results of the most recent PEAKS Studio 12.5 release (previously version 12.0). This led to 7,032 identified peptide sequences compared to 6,876 peptides with the older PEAKS version.^10^

#### FragPipe diaTracer coupled to DIA-NN

FragPipe (version 22.0) was used using the built-in “Nonspecific-HLA-diaPASEF” workflow outlined by Li *et al*.^23^ A target-decoy concatenated database was made using the proteome database described above. Briefly, diaTracer was used to deconvolute pseudo-MS/MS spectra from raw Bruker timsTOF (.d) runs. These spectra were searched using MSFragger (version 4.1)^25^ using the HLA-immunopeptide settings setting maximal peptide length to 20. Variable modifications included methionine oxidation and N-terminal protein acetylation (maximal one modification per peptide). Search results were re-scored using MSBooster (version 1.2.31),^26^ after which an empirical spectral library was made (library.tsv) that was searched by DIA-NN (version 2.0.2) in generic confidence scoring in ‘optimal results’ speed mode for quantification.^19^

#### Spectronaut directDIA

Spectronaut 19 (version 19.8.2503011.62635)^22^ directDIA was used for spectrum-centric analysis of the diaPASEF runs. The tims data (.tdf) were specified as input. For the Pulsar Search, the digest type was set to unspecific and maximal peptide length to 20. Variable modifications were set to N-terminal protein acetylation and methionine oxidation (maximal one modification per peptide). Protein FDR was set to 1 (100%) for identification and quantification. The default ‘Peptide Quant’ report was exported and used.

#### Multi-engine Rescoring Pipeline

Pseudo-MS/MS spectra (.mzML) outputted by diaTracer were subjected to a multi-engine rescoring pipeline described for the ddaPASEF analysis.^10^ Briefly, the pseudo-MS/MS spectra were searched by MSFragger^25^, Comet^27^, Sage^28^ and PEAKS Studio 12.5^29^ against a target-decoy concatenated database of the FASTA described above. Search engine results were rescored using TIMS^2^Rescore^30^ and results were aggregated at PSM and peptide-level as described for the ddaPASEF data.^10^ Results are filtered at 1% mokapot peptide Q-value and PSMs with the lowest posterior error probability (PEP) were used to extract retention time, ion mobility and matching b/y fragments to include in a spectral library for DIA-NN searching (version 2.0.2).^19^

### DIA-NN peptide-centric analysis using predicted libraries

#### Predicted spectral library generation

DIA-NN (version 2.0.2) was first used to predict spectral libraries encompassing all 8-to 12-mers of the human and *Listeria monocytogenes* strain EGD reference proteomes. To reduce RAM usage, we split the spectral libraries and first pass search per peptide precursor charge state, thus predicting 1+, 2+ and 3+ peptide precursor libraries with corresponding precursor *m/z* ranges (**Table 1**). Methionine oxidation was set as variable modification (maximal one modification per peptide).

**Table 1.**
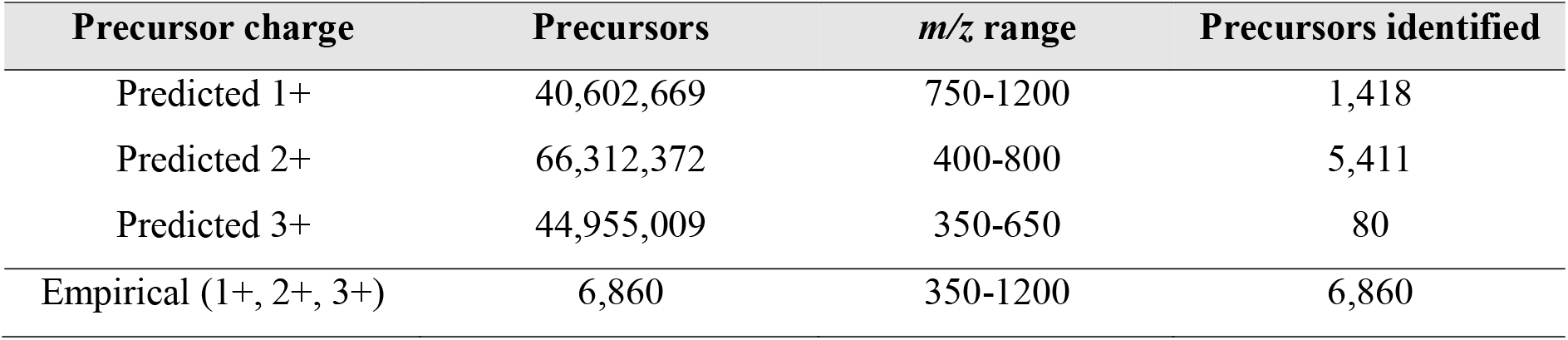
DIA-NN spectral library sizes and precursor identifications.

| Precursor charge | Precursors | *m/z* range | Precursors identified |
| --- | --- | --- | --- |
| Predicted 1+ | 40,602,669 | 750-1200 | 1,418 |
| Predicted 2+ | 66,312,372 | 400-800 | 5,411 |
| Predicted 3+ | 44,955,009 | 350-650 | 80 |
| Empirical (1+, 2+, 3+) | 6,860 | 350-1200 | 6,860 |

#### Spectral library searches

All raw diaPASEF (.d) data was searched with DIA-NN against each predicted charge-specific spectral library with proteoform confidence enabled and using the ‘low RAM and high speed’ mode. To further speed up data calibration in DIA-NN, we employed a small predicted spectral library of the peptide precursors included in the spectral library (library.tsv) generated by FragPipe on the diaTracer searches (using ‘*--ref*’) and additional options ‘*--rt-window-mul 1*.*7 -- rt-window-factor 100 --min-cal 500 --min-class 1000 --mass-acc-cal 25*’ that enable speed optimizations available in DIA-NN. All searches were conducted with a 15 ppm precursor and fragment tolerance (as recommended for timsTOF data). Automatic re-analysis (matching-between-runs, MBR) was disabled. Combining the results of three charge-specific searches, an empirical spectral library was generated by DIA-NN, incorporating 6,860 peptide precursors. In a final (second pass) search, this empirical library was used to analyze the same data in the generic confidence mode of DIA-NN, combined with the ‘optimal results’ speed mode.

##### Immunoinformatics and visualization

A standalone version of NetMHCpan-4.1b^31^ were used to predict the binding strength of immunopeptides to HeLa class I alleles HLA-A*68:02, HLA-B*15:03 and HLA-C*12:03 for HeLa. Alleles with the strongest predicted binding (lowest %Rank score) were used in the final reports, with a %Rank score below 0.5 and 2 are defined as strong and weak HLA-binding peptides, respectively. Unsupervised alignment and clustering of all 8-to 12-mer peptides was performed using GibbsCluster 2.0^32^ using recommended settings for MHC class I immunopeptides (motif length 9, max deletion length 4, max insertion length 1). Data organization and visualization was performed with custom Python scripts using pandas,^33^ matplotlib,^34^ and seaborn.^35^

## RESULTS AND DISCUSSION

### diaPASEF spectrum-centric analysis complements ddaPASEF analysis by detection of lower abundant immunopeptides

To evaluate DIA acquisition for bacterial immunopeptidomics, we analyzed purified immunopeptides from four uninfected and four *Listeria*-infected HeLa cell cultures in diaPASEF mode on a timsTOF SCP instrument (**Figure 1A**). These samples were previously analysed by ddaPASEF acquisition.^36^ An optimized diaPASEF window scheme was generated using the py_diAID package (**Figure 1B, Supplemental Table 1**).^24^ We first performed spectrum-centric analysis of the acquired diaPASEF data using FragPipe’s diaTracer^23^ (FP-diaTracer workflow) and Spectronaut’s directDIA^22^ algorithms (SN-dDIA workflow, **Figure 1C**). Both of these workflows deconvolute pseudo-MS/MS spectra from diaPASEF data, which can be searched by conventional DDA search engines. To replicate closer the ddaPASEF data that was analyzed with a multi-engine approach (combining MSFragger,^25^ Comet,^27^ Sage,^28^ and PEAKS Studio 12.5^29^) followed by rescoring with TIMS^2^Rescore^30^ (multi-engine DDA workflow)^10^, we also searched the diaTracer pseudo-MS/MS spectra with the same four search engines and rescoring approach (multi-engine DIA workflow). Both the FP-diaTracer and SN-dDIA workflows identified approximately 6,000 peptide sequences (**Dataset S1A-B**), a thousand peptides less compared to the multi-engine DDA workflow that identified 7,032 peptides (**Dataset S2, Figure 1D**). However, the multi-engine DIA workflow resulted in a total of 7,398 peptides (**Dataset S1C**), surpassing the multi-engine DDA workflow by 366 peptides (+5.2%). TIMS^2^Rescore boosted peptide identification in pseudo-MS/MS spectra over 60% for Comet, Sage and MSFragger, with the timsTOF model showing a 0.85–0.86 spectral Pearson correlation (**Supplemental Figure 1A-B**). Similarly, such rescoring is performed in the FragPipe workflow by MSBooster^26^ by the unweighted spectral entropy scoring feature, which also performs good on these pseudo-MS/MS spectra (**Supplemental Figure 1C**).

**Figure 1.**
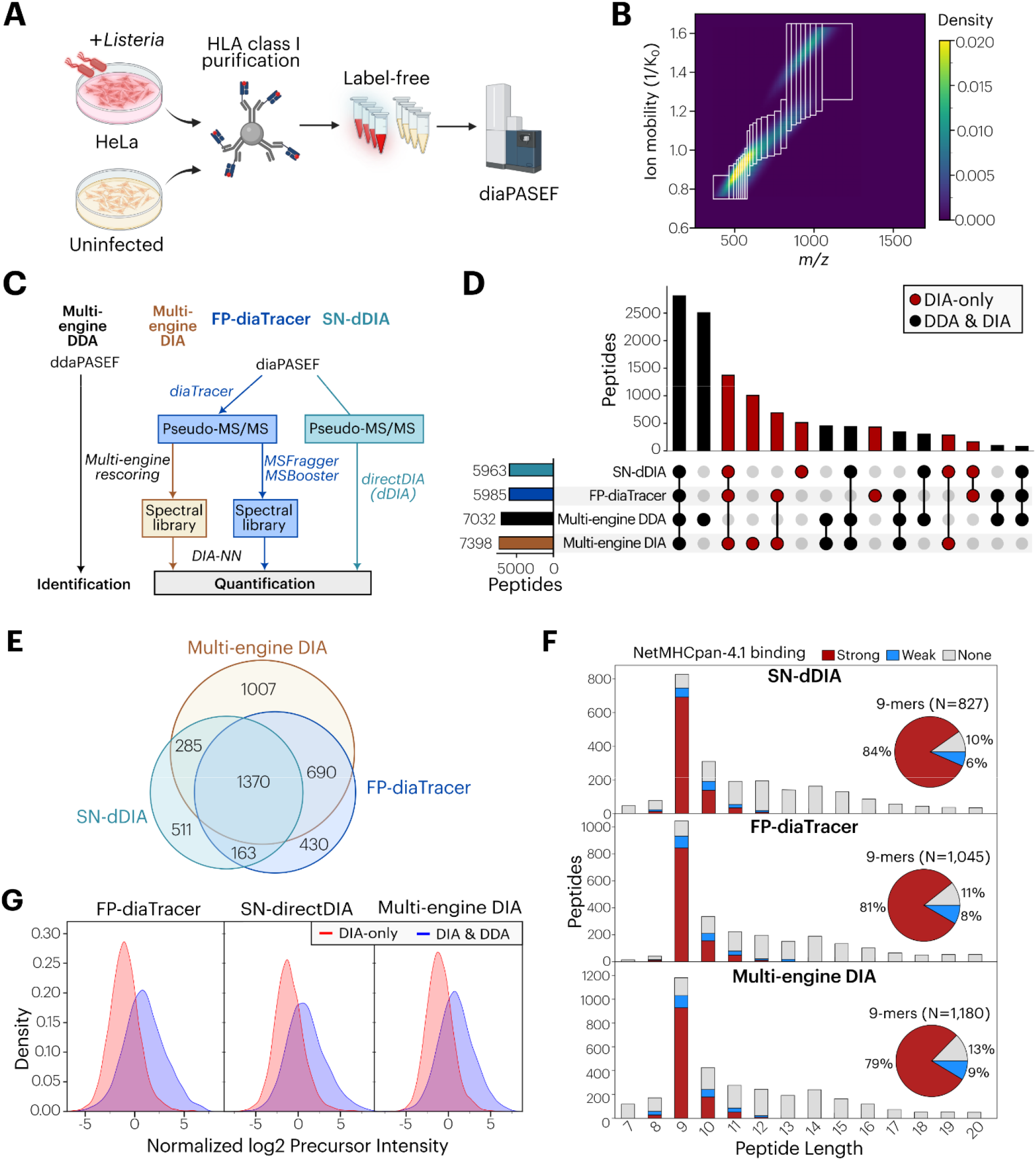
Spectrum-centric diaPASEF analysis complements ddaPASEF immunopeptide identifications. (**A**) Experimental set-up. HLA class I purified peptides of *Listeria*-infected and uninfected HeLa cells, previously analysed by ddaPASEF,^10^ were subjected to diaPASEF on a timsTOF SCP instrument. Created with BioRender.com. (**B**) Used diaPASEF acquisition scheme plotted on a kernel density distribution of all ddaPASEF identified precursors by py_diAID.^24^ Precursors were from a HLA class I immunopeptidomics sample of JY cells that is used as an in-house quality control. (**C**) Spectrum-centric analysis workflow. Raw Bruker diaPASEF (.d) files were converted to pseudo-MS/MS spectra via Spectronaut 19 directDIA or diaTracer. In case of Spectronaut directDIA (SN-directDIA), the built-in Pulsar search engine is used. In the FragPipe diaTracer analysis (FP-diaTracer), diaTracer pseudo-MS/MS spectra are searched by MSFragger results and rescored by MSBooster^26^ to generate a spectral library for DIA-NN quantification as described.^23^ Lastly, we applied our multi-engine rescoring pipeline^10^ to the diaTracer pseudo-MS/MS (mzML) files. Rescored results of all four search engines was used to generate a spectral library for DIA-NN quantification. (**D**) UpSet plot of the peptide sequences identified by each spectrum-centric approach and the ddaPASEF analysis. DIA-only peptide intersections were indicated in red, whereas those also identified by ddaPASEF are indicated in black. (**E**) Proportional Venn diagram of DIA-only identified peptides by spectrum-centric approaches. (**F**) Number of unique peptide sequences identified per amino acid length that were identified by spectrum-centric diaPASEF analysis but not by ddaPASEF analysis. Peptide sequences predicted as strong binder (SB, %Rank <0.5) or weak binder (WB, %Rank <2) by NetMHCpan-4.1^31^ are indicated in red and blue, respectively. Other peptides (nonbinder, NB) are indicated in gray. A piechart distribution of the DIA-only 9-mer peptide binding prediction was shown. (**G**) Density plots of all median normalized log2 precursor intensities across all samples measured by SN-dDIA, FP-diaTracer and multi-engine DIA workflows. Density plots were shown for precursors of peptide sequences also identified by ddaPASEF (blue) or uniquely in diaPASEF analysis (red).

A total of 2,812 peptides were identified in all four workflows (**Figure 1D**), while 2,507 peptide sequences were identified solely by the multi-engine DDA workflow and 4,456 peptides were unique to at least one of the spectrum-centric DIA workflows (**Figure 1E**). To assess the quality of these 4,456 DIA-only immunopeptides, we inspected their peptide length distribution and predicted binding capacity to HeLa MHC class I alleles (HLA-A*02:01, HLA-B*07:02 and HLA-C*07:02) using NetMHCpan-4.1.^31^ For all three workflows the DIA-only peptides show a dominant 9-mer peak consisting of 87-90% of predicted HLA binders (**Figure 1F**), thus indicating high-quality immunopeptide identifications, with the multi-engine DIA workflow resulting in the highest identification numbers, followed by FP-diaTracer. Interestingly, when evaluating the peptide precursor intensity, we observed that for each of the DIA workflows the DIA-only peptides were significantly lower intense compared to peptides also identified by ddaPASEF (**Figure 2C**). This observation was also supported by testing the reverse scenario where the ddaPASEF data was re-analyzed quantitatively with FragPipe enabling IonQuant^37^ for MS1-based quantification. Here, DDA-only peptides not identified by diaPASEF were equally abundant as peptides also identified by diaPASEF (**Supplemental Figure 2**). Together, these data show that diaPASEF acquisition of immunopeptides in combination with spectrum-centric data analysis complements ddaPASEF analysis, detecting additional lower abundant immunopeptides.

### diaPASEF peptide-centric analysis recovers low abundant immunopeptides

Non-enzymatic proteome searches are computationally complex given the millions of peptide possibilities to be searched. To reduce this complexity for DIA immunopeptidomics searches, previous studies have generated *in silico* spectral libraries restricted to peptides predicted to bind the HLA alleles of the biological sample.^12,13^ In this study we aimed to achieve maximum sensitivity of HLA peptide detection using unrestricted peptide-centric searches of predicted spectral libraries encompassing all possible MHC class I peptides (length 8-12) of the combined human and *Listeria* reference proteomes. To this end, charge-specific (1+, 2+, and 3+) spectral libraries of all 8 to 12 amino acids long peptides were predicted using DIA-NN (allowing a single methionine oxidation), together spanning ∼152 million peptide precursors (**Figure 2A**). These were searched against the data, resulting in three per-charge DIA-based empirical spectral libraries that were combined. Subsequently, searching the combined library yielded a total of 6,845 precursors corresponding to 6,535 peptide sequences (**Dataset S3**). Identified HLA class I peptides were of high-quality, demonstrated by the identification of 4,073 9-mers including 92% predicted HLA binders (**Figure 2B, Supplemental Figure 3A**).

**Figure 2.**
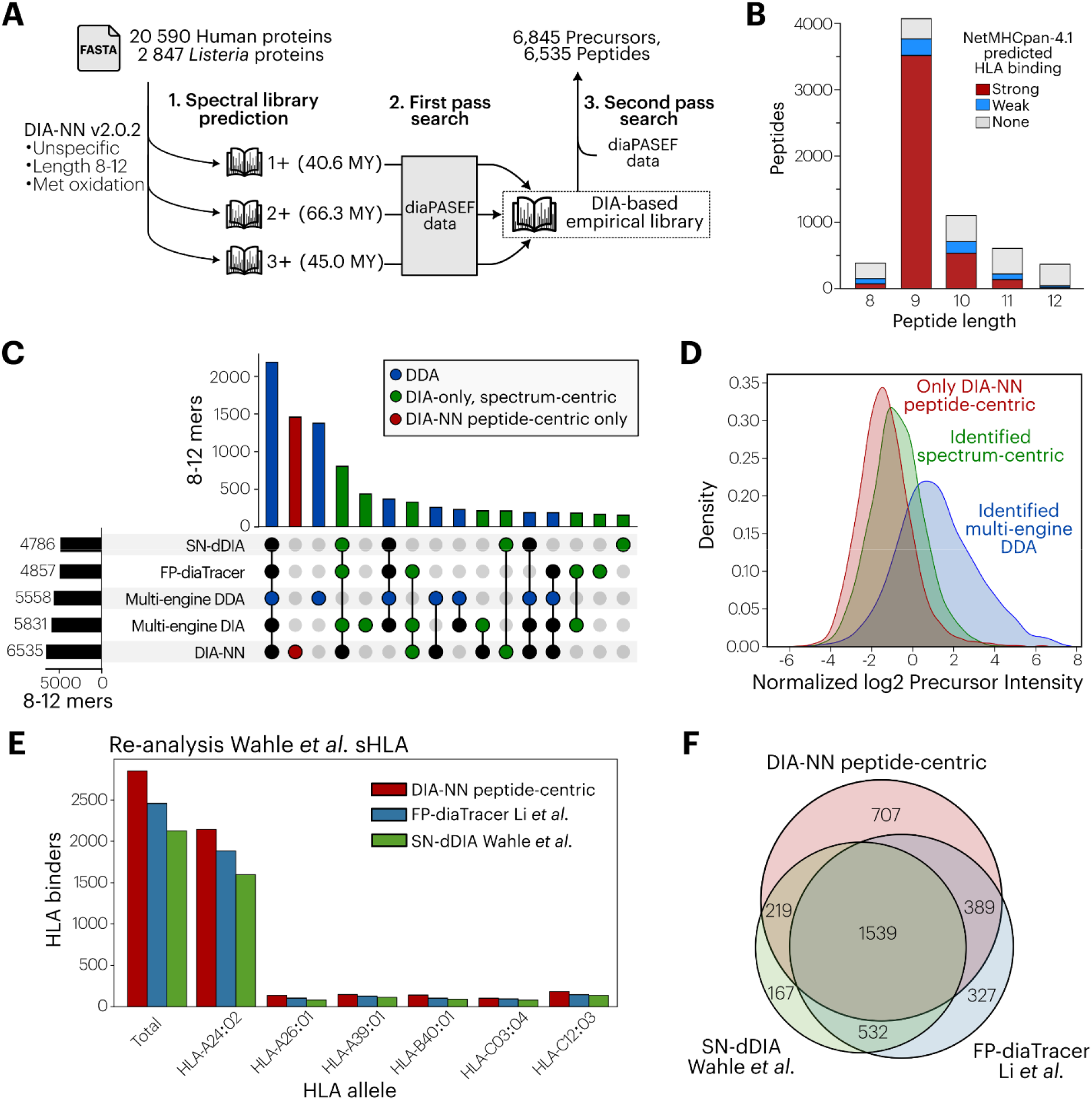
Unspecific peptide-centric immunopeptidomics searches with DIA-NN identify lower-abundant peptide precursors. (**A**) Peptide-centric search strategy. Step 1: human and *Listeria monocytogenes* EGD UniProtKB reference proteomes were used for generating predicted spectral libraries of all 8-12 mers per charge state (1+, 2+ or 3+). Step 2: these predicted libraries are searched (first pass) in proteoform scoring mode. Step 3: resulting empirical DIA-based libraries are searched collectively in a second pass search, identifying a total of 6,845 precursors matching 6,535 8-12 mer peptides. (**B**) Number of unique peptide sequences (N=6,535) identified per amino acid length that wer identified by peptide-centric DIA-NN analysis. Peptide sequences predicted as strong binder (SB, %Rank <0.5) or weak binder (WB, %Rank ≤ 2) by NetMHCpan-4.1^31^ are indicated in red and blue, respectively. Other peptides (nonbinder, NB) are indicated in gray. (**C**) UpSet plot of the peptide sequences identified by the peptide-centric DIA-NN search, the spectrum-centric approaches and the ddaPASEF analysis. Peptides identified by ddaPASEF were indicated in blue, DIA-only peptides found by any spectrum-centric analysis in green, and peptides unique to DIA-only peptide-centric analysis by DIA-NN in red. (**D**) Density plots of log2 precursor QuantUMS intensities measured by DIA-NN workflow for precursors of peptide sequences identified by ddaPASEF (blue), DIA-only spectrum-centric analyses (green) or uniquely by DIA-NN peptide-centric analysis (green). (**E**) Identified HLA binders (%Rank ≤ 2) from re-analysis of a healthy donor plasma sample from the study of Wahle *et al*.^12^ (using Spectonaut 17 directDIA) that was also re-analysed by diaTracer.^23^ (**F**) Proportional Venn diagram of identified HLA binders (%Rank ≤ 2) identified by DIA-NN predicted immunopeptide library searches in this work and reported by SN-dDIA by Wahle *et al*.^12^ and diaTracer by Li *et al*.^23^

The peptide-centric searches of DIA-NN using the predicted libraries outperformed the spectrum-based analyses in terms of quantified 8-12 mer peptide sequences. It identified more 8-12 mers than the SN-dDIA and FP-diaTracer workflows (34-36% increase) as well as the multi-engine DDA and DIA workflows (12-18% increase) (**Figure 2C**). Moreover, there was a large proportion of 1,458 immunopeptides that were uniquely identified by DIA-NN (**Figure 2C, Supplemental Figure S4**). These DIA-NN-only peptides were of high-quality with a dominant peak of 9-mers (783 peptides) including 85% predicted HLA binders (**Supplemental Figure 3B**). In terms of intensity, precursors of these DIA-NN-only peptides were yet lower abundant than those identified in the spectrum-centric analyses (**Figure 2D**). This suggests that these very low-abundant immunopeptides are generally missed in spectrum-centric analyses, possibly due their low intensity signals challenging confident pseudo-MS/MS generation. However, direct interrogation of diaPASEF data with spectral libraries levering deep learning-predicted peptide features does seem to distinguish these low-abundant precursors, in line with previous observations.^38^

To test whether the increased performance of these immunopeptide searches using proteome-wide predicted libraries generalizes to other datasets we re-analyzed three diaPASEF plasma immunopeptidome samples that were analyzed by Wahle *et al*.^12^ using Spectronaut 17 dDIA. Interestingly, these samples were already re-analyzed by Li *et al*.^23^ using FP-diaTracer. We searched ∼140 MY human peptide precursors of 8 to 12 amino acids long with our peptide-centric DIA-NN workflow. In total, we identified 2,975 peptide sequences that included 2,851 (96%) predicted binders (%Rank ≤ 2) to the respective donor HLA alleles (**Dataset S4**). This presented an additional 394 and 721 predicted binders (16% and 34% increase) compared to those reported by Li *et al*.^23^ and Wahle *et al*.^12^, respectively, showing a similar predominance of HLA-A*24:01 peptide binders (**Figure 2E**). These DIA workflows showed complementary identifications (**Figure 2F**) with 707 HLA binders unique to the DIA-NN searches that were lower abundant than those identified by either SN-dDIA or FP-diaTracer (**Supplemental Figure 5A**). The reliability of these DIA-NN-only peptides is supported by the fact the majority – 566 out of 707 peptides (80%) – was also identified by the DDA and pan spectral library searches of Wahle *et al*.^12^ (**Supplemental Figure 5B**). Together, these results highlight the added value of peptide-centric searches for proteome-wide discovery of immunopeptides, particularly for detection of low-abundant immunopeptides.

### diaPASEF analyses uncover low-abundant *Listeria* immunopeptides

After demonstrating the merit of both peptide- and spectrum-centric searches for global immunopeptide identification, we next focused on their capacity to detect and quantify *Listeria*-derived immunopeptides presented on the infected HeLa cells (**Figure 1A**). To this end, we filtered high confident *Listeria* peptide precursors from all DIA search results. Firstly, we verified that *Listeria* peptide sequences did not match human proteins, common contaminants or the nuORF database^39^ (considering Ile equal to Leu). Solely the 7-mer peptide ‘VSLLKEI’ was discarded, being isobaric to the identified ‘VSIIKEL’ matching multiple human proteins. Secondly, we required *Listeria* peptide precursors to be quantified in at least two replicate samples of the infected condition while being absent in all replicates of the uninfected condition. The majority (70-85%) of peptides fulfilled these criteria over the various DIA analysis workflows, with the DIA-NN peptide-centric workflow showing the least quantified signal in the mock condition (**Supplemental Figure 6**). Across all DIA workflows, a total of 37 bacterial immunopeptides matching 27 *Listeria* proteins fulfilled these criteria, often quantified in three or four infected replicate samples (**Dataset S5-6, Figure 3A**). Eleven *Listeria* peptides were identified by all DIA analysis workflows, seven of which were also picked up in the ddaPASEF (multi-engine DDA) analysis (**Figure 3B**). Peptide-centric analysis by DIA-NN recovered a relatively high number of ten *Listeria* peptides not found by any spectrum-centric analysis, seven of which were also not found in the ddaPASEF analysis (**Figure 3B**). The high confidence of the identified *Listeria* peptides is further supported by the fact that 34 out of 37 peptides (92%) were predicted binders to HeLa HLA class I alleles (**Figure 3C**). This number exceeds the 27 out of 33 (81%) *Listeria* peptides that we previously identified as predicted binders by ddaPASEF.^10^ In addition, these 37 high confident bacterial peptides match several *Listeria* antigens that were previously described and/or identified by ddaPASEF analysis.^4,10^ diaPASEF analysis did identify an additional peptide for the effector protein phospholipase C (plcB) and the peptidoglycan hydrolase LMON_2714, as well as detected predicted binders for yet unidentified *Listeria* proteins including cysK, OppD, rpoD, rplF and others (**Figure 3C**).

**Figure 3.**
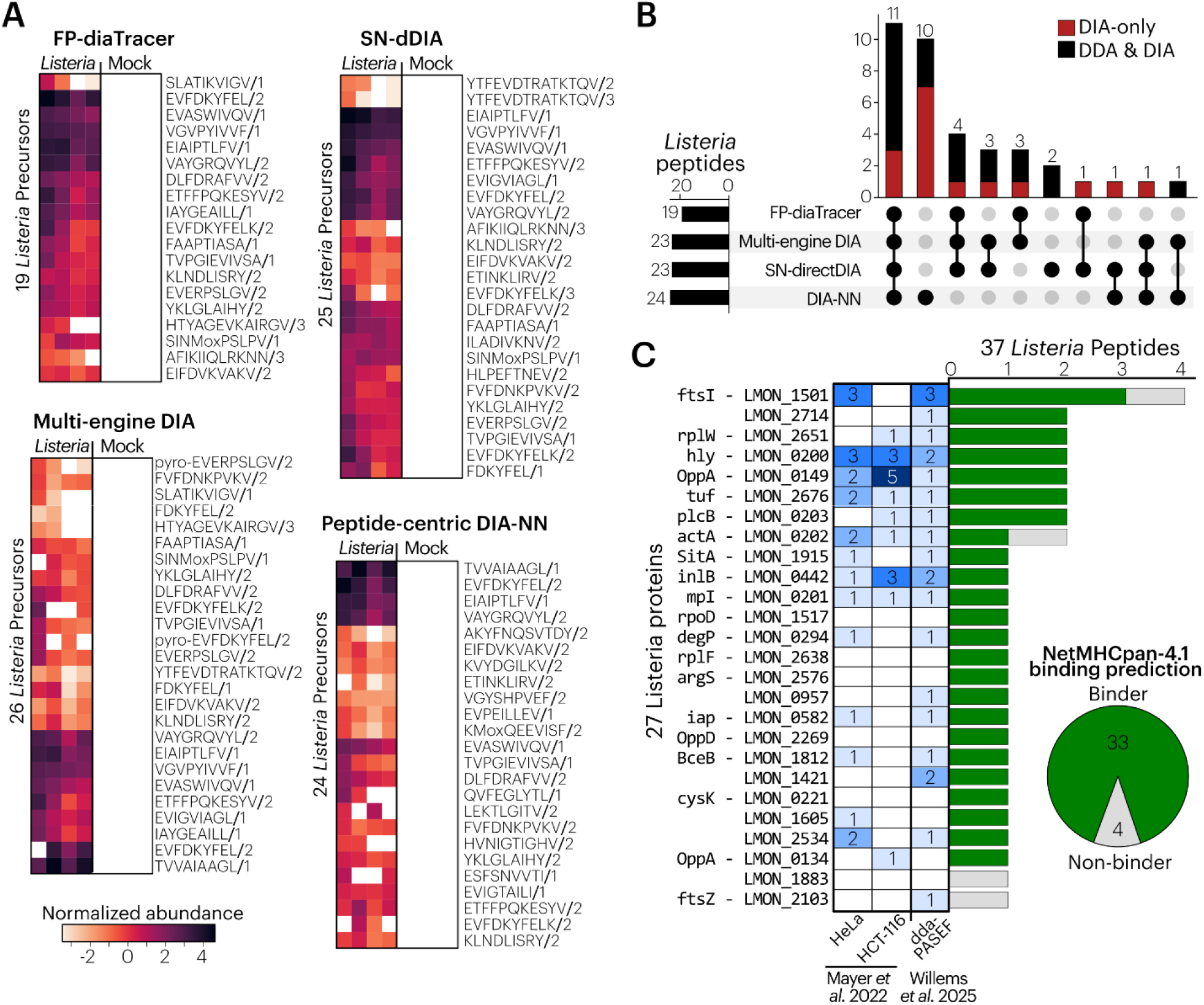
Bacterial epitope discovery by data-independent acquisition immunopeptidomics. (**A**) Heatmaps of identified *Listeria* peptide precursors log2 displaying the normalized intensities outputted b each DIA analysis workflow. High-confidence *Listeria* peptides were filtered for not matching any human, contaminant or nuORFdb^39^ proteins (considering Ile equal to Leu) and were required to be quantified at least twice in the infected condition and absent in the non-infected condition. (**B**) UpSet plot of high-confidence *Listeria* peptides identified by DIA analyses. Peptides found in the corresponding ddaPASEF runs were indicated in black. (**C**) The number of unique immunopeptides identified per *Listeria* is shown in a histogram. Immunopeptides predicted as HLA class I binders by NetMHCpan-4.1^31^ (%rank < 2) are indicated in green and peptides note predicted as HLA binders in grey. A heatmap indicates the number of peptides identified by Mayer *et al*.^4^ and in the ddaPASEF acquisition^10^ for these *Listeria* proteins.

Next, we inspected the intensities of the high confident *Listeria* epitopes. As observed previously,^10^ *Listeria* peptides showed an overall similar intensity as human peptide (**Supplemental Figure S7**). Thus, detected *Listeria* peptides are not lower abundant in terms of recorded MS intensity, though they are outnumbered by thousands of human self-peptides. Interestingly however, *Listeria* peptides only identified by diaPASEF were of lower abundance than those also found by ddaPASEF (**Figure 4A**). Furthermore, the seven *Listeria* peptides solely identified by the DIA-NN peptide-centric workflow tend to be of very low abundance. Illustrating this, the abundant *Listeria* peptide precursor ‘EVFDKYFEL/2’ matching FtsI was identified in every DIA and DDA analysis, and it chromatogram shows the ms1 precursor signal reaching up to ∼1.5 x 10^5^ intensity (**Figure 4B**). Conversely, the *Listeria* precursor ‘HVNIGTIGHV/2’ was only recovered peptide-centric workflow with a ∼3,7 x 10^3^ intensity and fragment ion intensities closer to the limit of detection (**Figure 4C**). Interestingly, we previously identified ‘HVNIGTIGHV’, a predicted HLA-A*68:02 binder, in TMT-labeled counterparts of the current HeLa samples analyzed on a Fusion Lumos mass spectrometer, while the peptide was missed in the label-free analysis.^4^ Together, these results demonstrate how peptide-centric DIA analysis aid in the detection of low-abundant bacterial immunopeptides.

**Figure 4.**
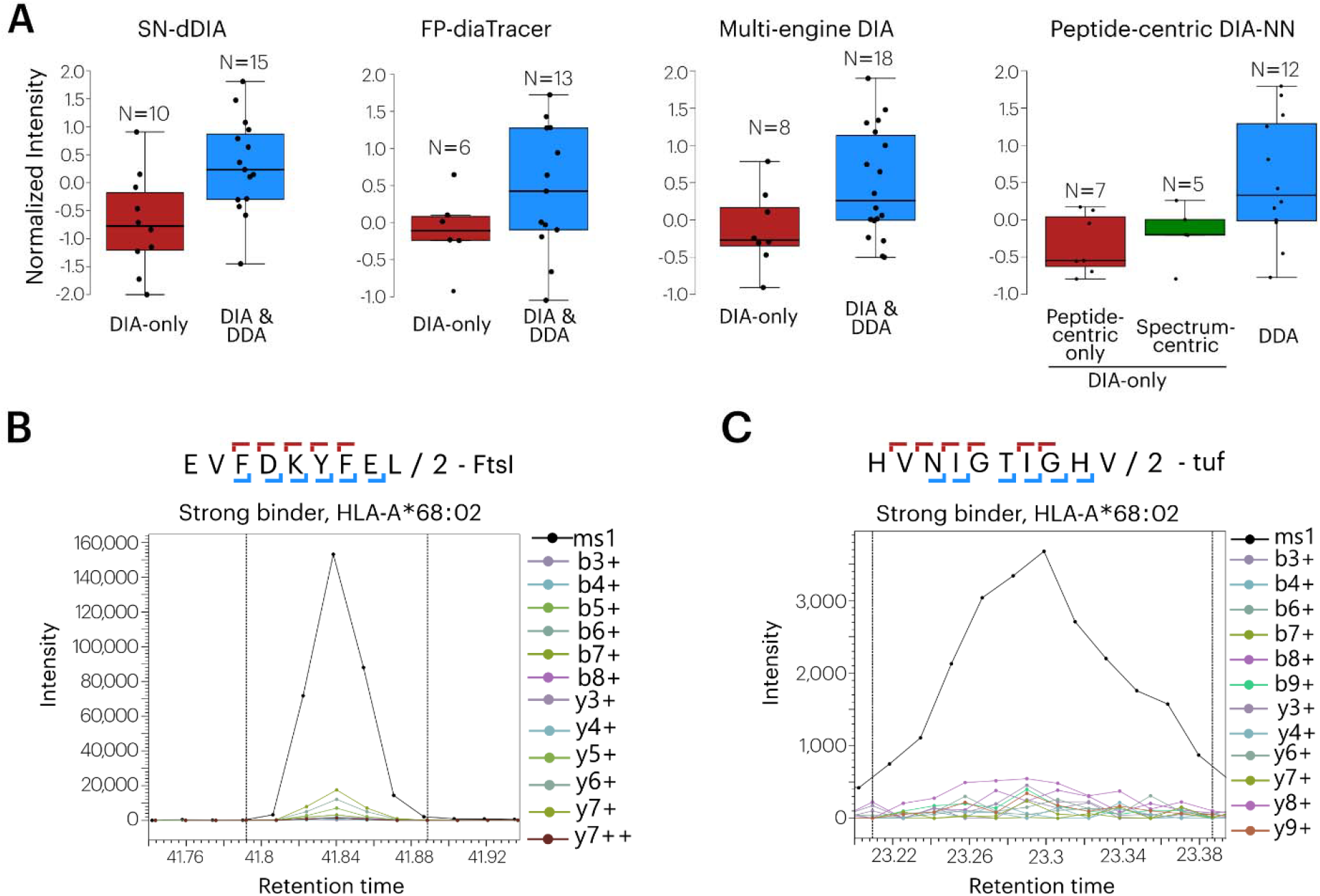
Spectrum-centric and peptide-centric DIA analyses recover low-abundant bacterial immunopeptides. (**A**) Normalized intensity of *Listeria* peptides for DIA-only peptides and those also identified by the corresponding ddaPASEF runs per DIA search strategy. In case of peptide-centric DIA-NN searches, we further distinguished peptides only identified by the peptide-centric search in DIA-NN using predicted libraries. (**B-C**) Extracted chromatograms by DIA-NN Viewer for EVFDKYFEL/2+ (**B**) and HVNIGTIGHV/2+ (**C**). Ms1 precursor signal is indicated in black, while fragment ion in a color palette (see legend). Peak boundaries are indicated as dotted vertical lines. For chromatograms of all high-quality *Listeria* peptides see Supplemental Data S6.

## CONCLUSIONS

DIA analysis is increasingly adopted in immunopeptidomics to boost comprehensive detection and quantification of immunopeptides.^12,14-16^ In this work, we demonstrate how spectrum- and peptide-centric analysis of diaPASEF data shows great promise to detect complementary lower-abundant identifications missed by DDA. Whereas the FP-diaTracer and SN-dDIA workflow offer proteome-wide unspecific searches of pseudo-MS/MS spectra, we demonstrate spectral library prediction and searching with DIA-NN of all 8-12 mer peptides to be possible and recover a higher number of identified MHC class I peptides. Likely, increased identification could still be gained by fine-tuning deep learning models for peptide properties prediction to timsTOF fragmentation and/or DIA acquisition. For instance, Carafe trains its fragment ion intensity and retention time prediction model directly on the DIA MS/MS spectra itself, thereby boosting peptide identification.^40^ The performance and feasibility of these approaches is bound to increase with experiment-tuned deep learning models and the ever-continuing advances in computational power. While we showcase how DIA immunopeptidomics can be strengthened, greatest depth can be achieved by combining DDA and DIA injections. This allows to maximize the detection of bacterial immunopeptides as an emerging approach for antigen discovery in bacterial vaccine development.^1^

## ASSOCIATED CONTENT

### Data availability statement

The diaPASEF mass spectrometry proteomics data have been deposited to the ProteomeXchange Consortium (http://proteomecentral.proteomexchange.org) via the PRIDE partner repository^41^ with the dataset identifier PXD063560. Previous recorded ddaPASEF acquired data from these samples is available via the dataset identifier PXD055547. Search results and code used for and visualization are available at https://github.com/diaPASEF_immunopeptidomics.

## Supporting Information

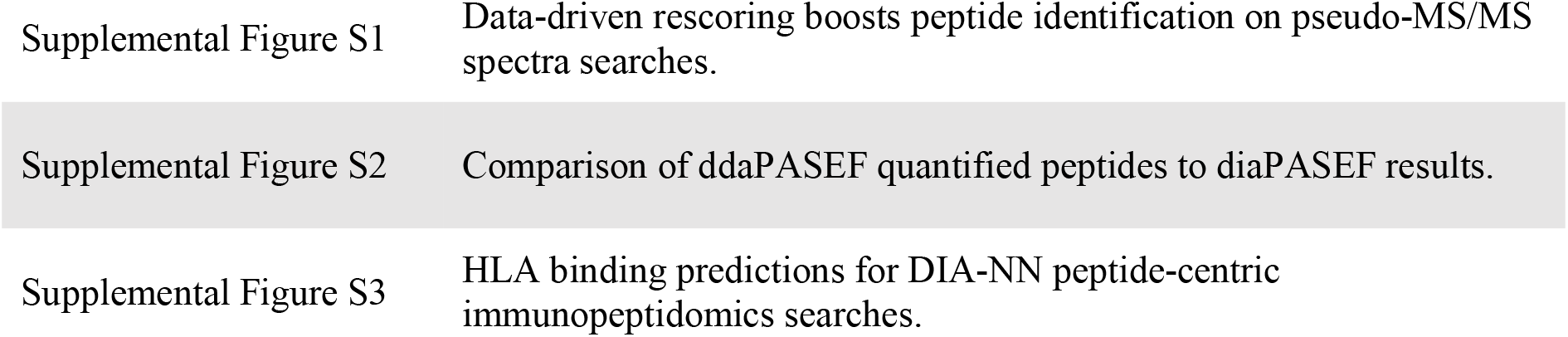
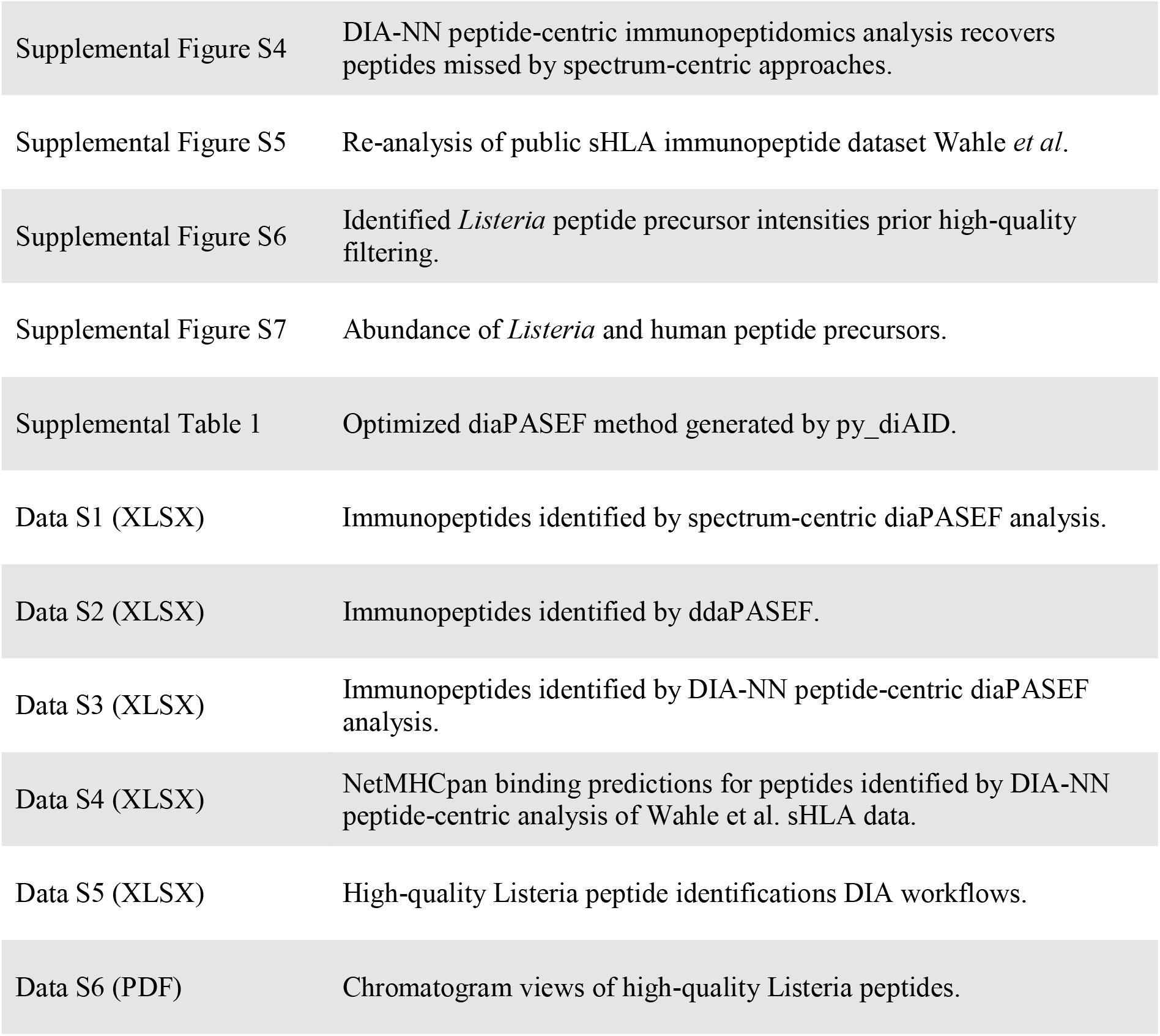

## AUTHOR INFORMATION

## Author Contributions

The manuscript was written through contributions of all authors. All authors have given approval to the final version of the manuscript.

## Notes

V.D. holds shares of Aptila Biotech. Other authors declare no competing financial interest.

## ACKNOWLEDGMENTS

We thank the VIB Proteomics Core for LC-MS/MS analysis. F.I. acknowledges support from Ghent University Concerted Research Action grant BOF21/GOA/033 and the Horizon Europe Project BAXERNA 2.0 (101080544). V.D. is supported by the German Ministry of Education and Research (BMBF), as part of the National Research Node “Mass spectrometry in Systems Medicine” (MSCoreSys), under grant agreement 161L0221.

## ABBREVIATIONS

DDA: data-dependent acquisition
DIA: data-independent acquisition
NB: non-binder
SB: strong binder
WB: weak binder

