## Supporting Information for "Data-independent immunopeptidomics discovery of low-abundant bacterial epitopes"

**TABLE OF CONTENTS**

| Supplemental Figure S1 | Data-driven rescoring boosts peptide identification on pseudo-MS/MS spectra searches. |
| --- | --- |
| Supplemental Figure S2 | Comparison of ddaPASEF quantified peptides to diaPASEF results. |
| Supplemental Figure S3 | HLA binding predictions for DIA-NN peptide-centric immunopeptidomics searches. |
| Supplemental Figure S4 | DIA-NN peptide-centric immunopeptidomics analysis recovers peptides missed by spectrum-centric approaches. |
| Supplemental Figure S5 | Re-analysis of public sHLA immunopeptide dataset Wahle *et al*. |
| Supplemental Figure S6 | Identified *Listeria* peptide precursor intensities prior high-quality filtering. |
| Supplemental Figure S7 | Abundance of *Listeria* and human peptide precursors. |
| Supplemental Table 1 | Optimized diaPASEF method generated by py_diAID. |
| Data S1 (XLSX) | Immunopeptides identified by spectrum-centric diaPASEF analysis. |
| Data S2 (XLSX) | Immunopeptides identified by ddaPASEF. |
| Data S3 (XLSX) | Immunopeptides identified by DIA-NN peptide-centric diaPASEF analysis. |
| Data S4 (XLSX) | NetMHCpan binding predictions for peptides identified by DIA-NN peptide-centric analysis of Wahle et al. sHLA data. |
| Data S5 (XLSX) | High-quality Listeria peptide identifications DIA workflows. |
| Data S6 (PDF) | Chromatogram views of high-quality Listeria peptides. |

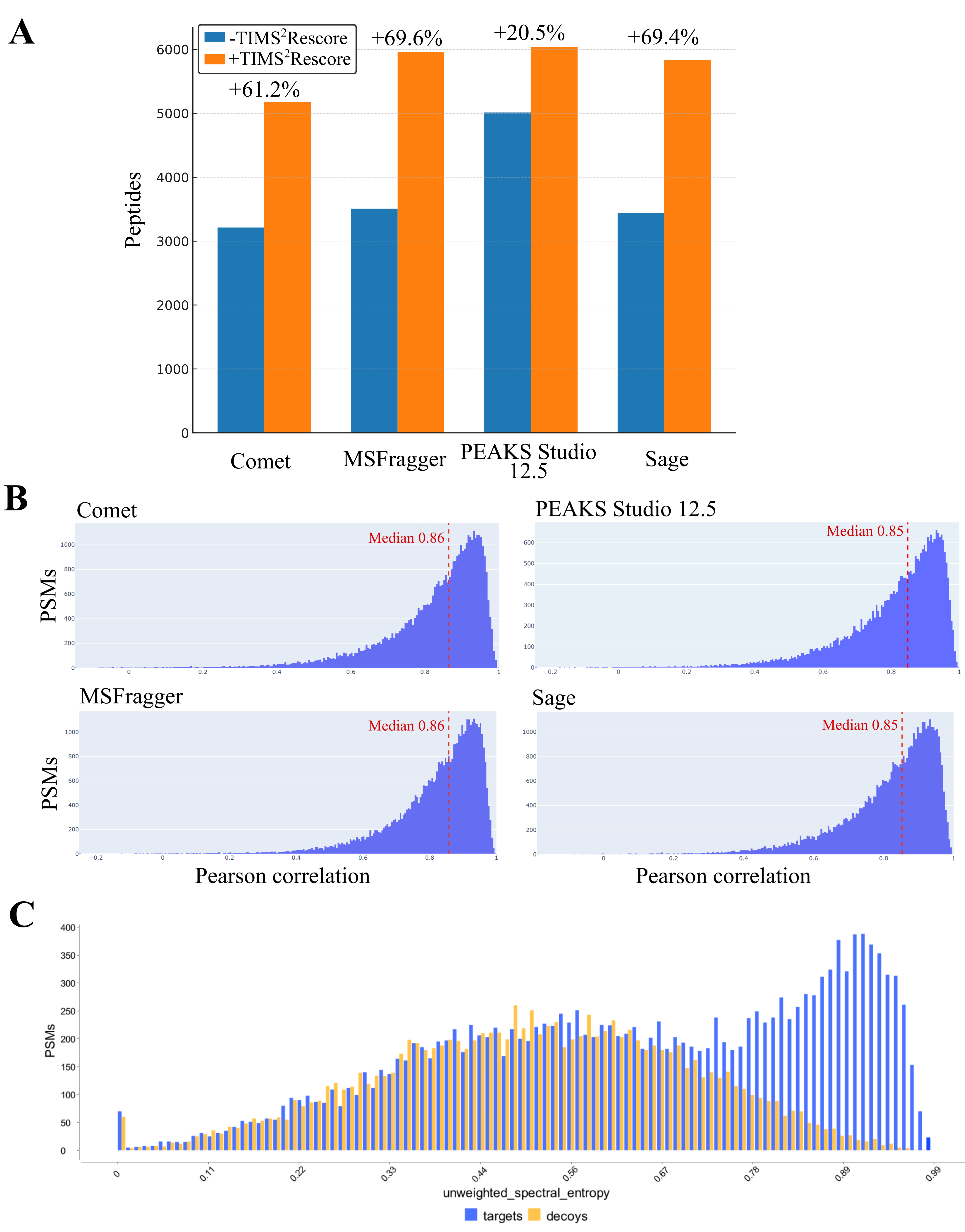

**Supplemental Figure S1. Data-driven rescoring boosts peptide identification on pseudo-MS/MS spectra searches in the multi-engine DIA workflow.** (**A**) Peptides identified before and after TIMS^2^Rescore^1^ rescoring (1% peptide FDR) per search engine when searching diaTracer^2^ pseudo-MS/MS spectra. Numbers are extracted from TIMS^2^Rescore generated reports. (**B**) Pearson correlation histograms of peptide-to-spectrum matches (PSMs) per search engine to MS/MS spectra predicted by the MS^2^PIP timsTOF model used by TIMS^2^Rescore. Vertical dotted lines indicate the median Pearson correlation. (**C**) Representative histogram of the unweighted spectral entropy score for target and decoy PSMs by MSBooster^3^ within the FP-diaTracer search. This was the fourth replicate of the Listeria-infected sample, but is representative for all other runs.

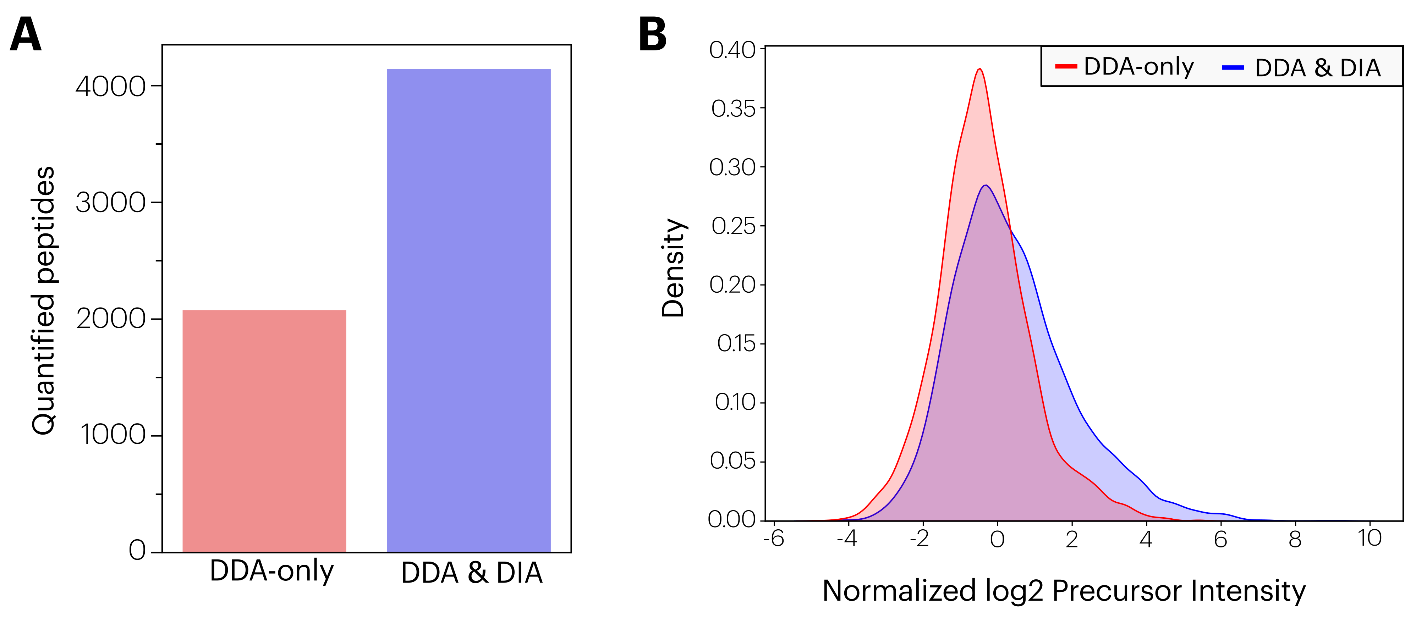

**Supplemental Figure S2. Comparison of ddaPASEF quantified peptides to diaPASEF results.** (**A**) Barchart of peptides identified by the FragPipe-HLA workflow. Peptides only identified by ddaPASEF were indicated in red, and those also present in the diaPASEF analysis were indicated in blue. (**B**) Density plots of sample median normalized peptide intensity measured by FragPipe IonQuant (derived from combined_peptide.tsv) for peptides solely identified by ddaPASEF (‘DDA-only’) or also identified in the diaPASEF spectrum-centric analyses.

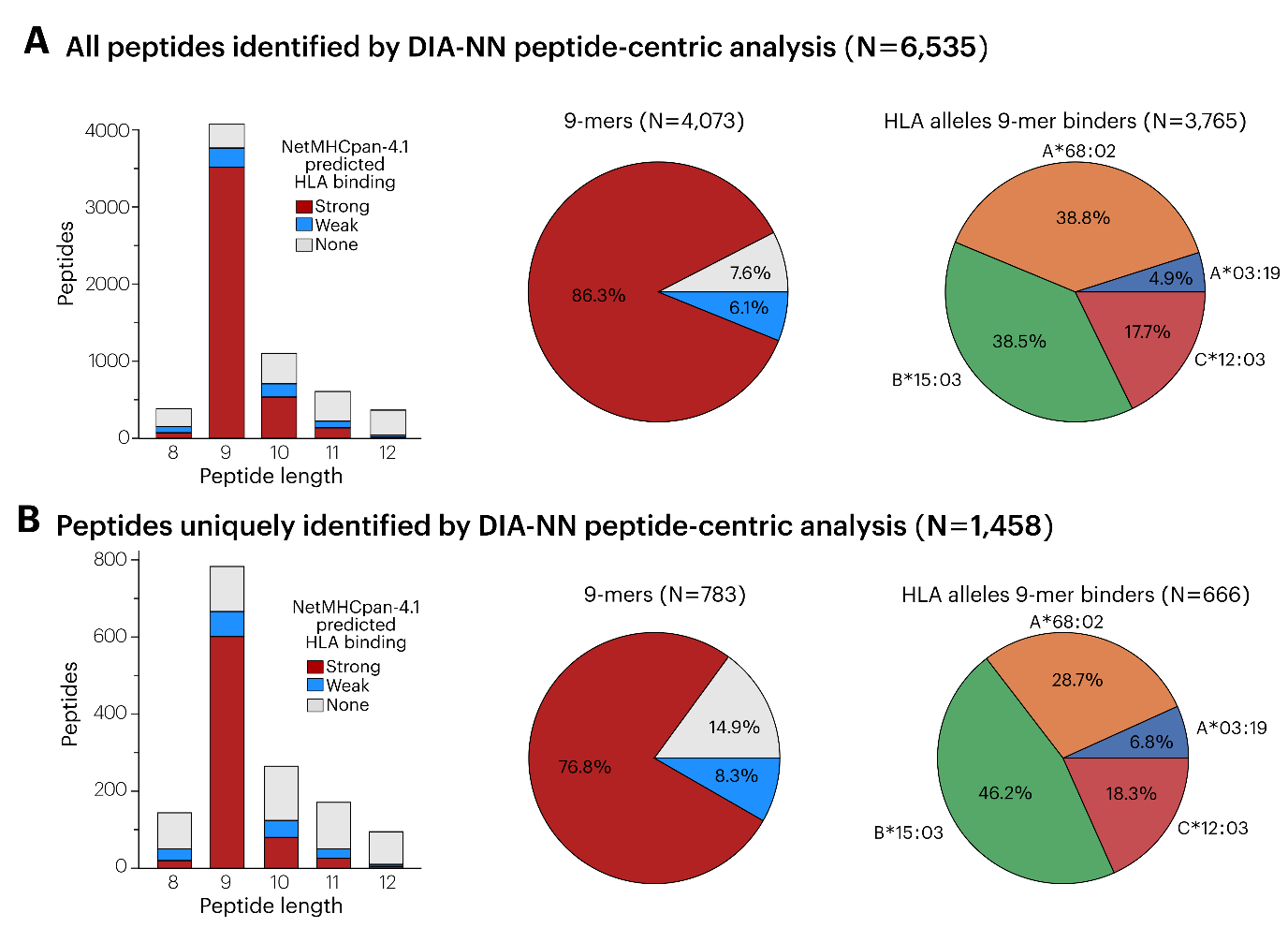

**Supplemental Figure S3. HLA binding predictions for DIA-NN peptide-centric immunopeptidomics searches.** HLA binding prediction was performed for all identified peptide sequences (**A**) and those unique to the DIA-NN peptide-centric analysis not found by ddaPASEF and diaPASEF spectrum-centric analysis (**B**). (*Left*) Number of unique peptide sequences (N=6,535) identified per amino acid length that were identified by peptide-centric DIA-NN analysis. Peptide sequences predicted as strong binder (SB, %Rank <0.5) or weak binder (WB, %Rank <2) by NetMHCpan-4.1^4^ are indicated in red and blue, respectively. Other peptides (nonbinder, NB) are indicated in gray. (*Right*) Pie chart distributions of HLA class I binding predictions. The HLA allele with the lowest %Rank score was plotted.

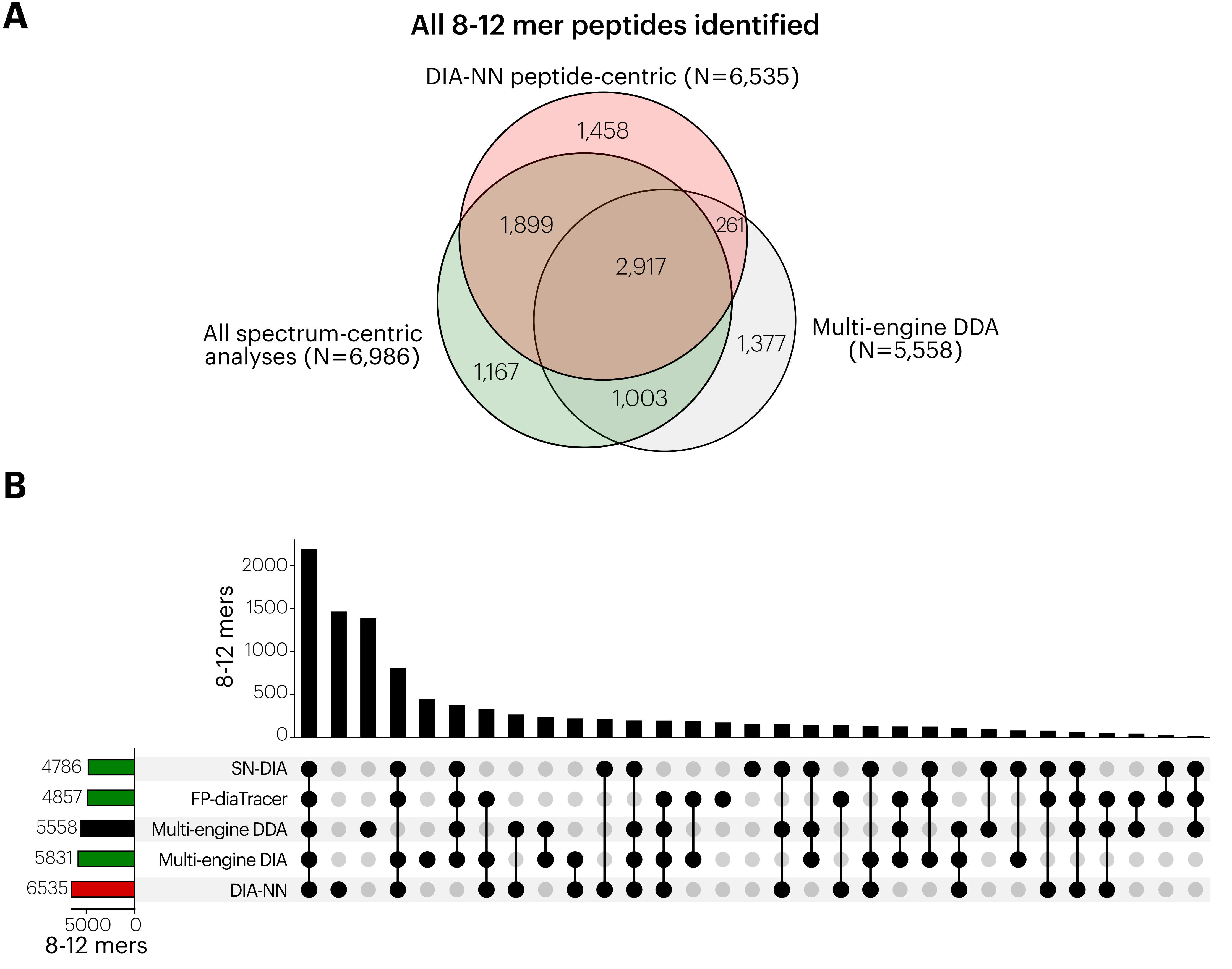

**Supplemental Figure S4. DIA-NN peptide-centric immunopeptidomics analysis recovers peptides missed by spectrum-centric approaches.** (**A**) Weighted Venn diagram of 8-12mer peptide sequences identified in the multi-engine ddaPASEF analysis, spectrum-centric diaPASEF analyses (SN-dDIA, FP-diaTracer or multi-engine DIA workflows indicated in green) and peptide-centric DIA-NN analysis (indicated in red). (**B**). UpSet plot of the 8-12 mer peptide sequences identified by all diaPASEF and the ddaPASEF workflows.

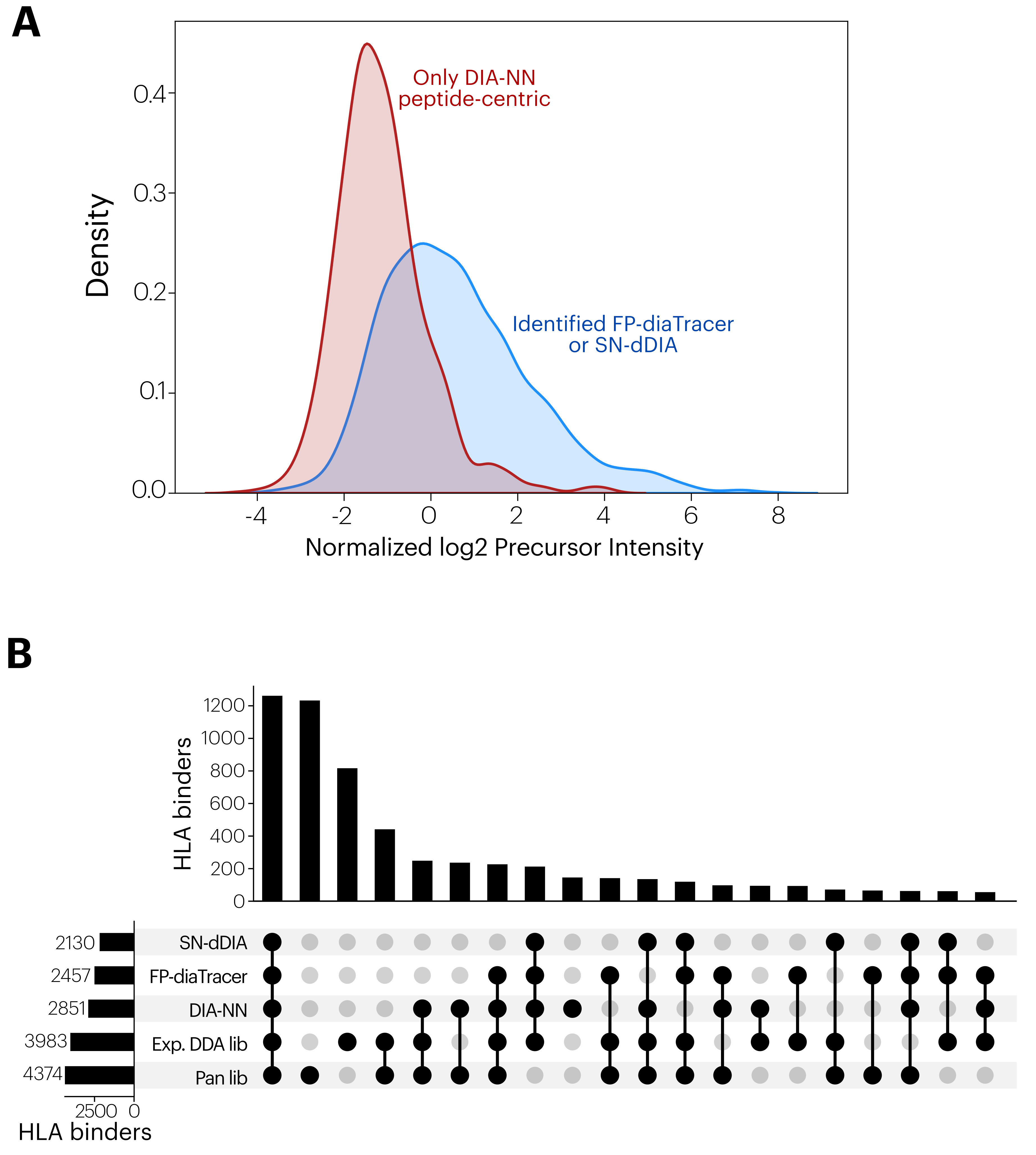

**Supplemental Figure S5. Re-analysis of public sHLA immunopeptide dataset Wahle *et al*.^5^** (**A**) Density plots of log2 precursor QuantUMS intensities measured by DIA-NN workflow for precursors of predicted HLA peptide binders identified by FP-diaTracer in Li *et al.*^2^ and/or SN-dDIA in Wahle *et al.*^5^ (blue) and the density plot of precursors uniquely identified by DIA-NN peptide-centric analysis (red). (**B**). UpSet plot of the predicted HLA binder (8-12 mers) peptide sequences identified by peptide-centric DIA-NN analysis, FP-diaTracer in Li *et al.*^2^, and SN-dDIA, experimental DDA library or pan library searches described in Wahle *et al.*^5^. Peptide intersections lower than 50 peptides were not plotted.

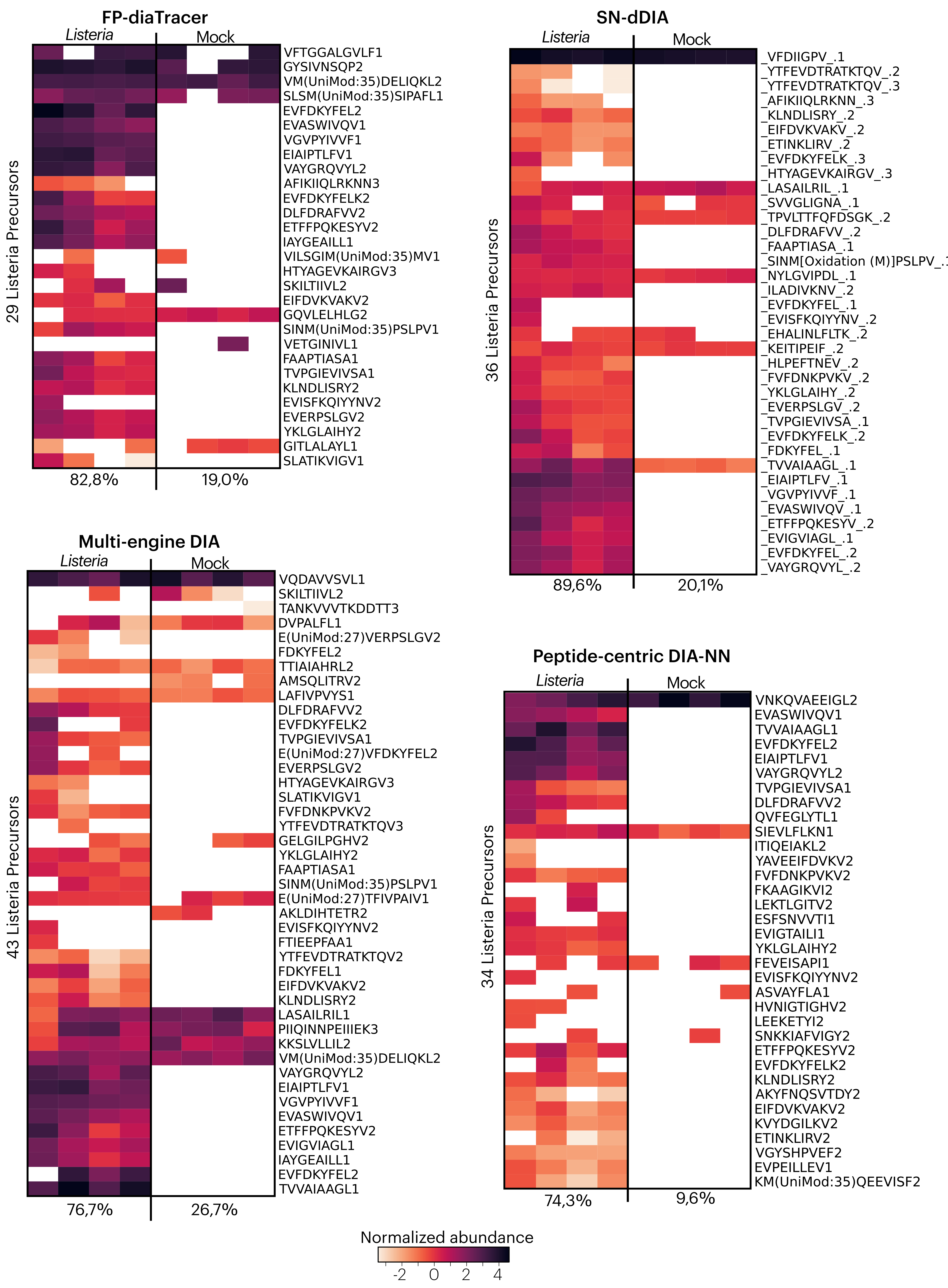

**Supplemental Figure 6. *Listeria* peptide quantification by data-independent acquisition immunopeptidomics.** Heatmaps of identified *Listeria* peptide precursors log2 displaying the normalized intensities outputted by each DIA analysis workflow. *Listeria* peptides were filtered for not matching any human, contaminant or nuORFdb^6^ proteins (considering Ile equal to Leu). Percentages below indicate the data completeness or proportion of valid quantified intensities for all peptides in the *Listeria*-infected and uninfected, mock condition.

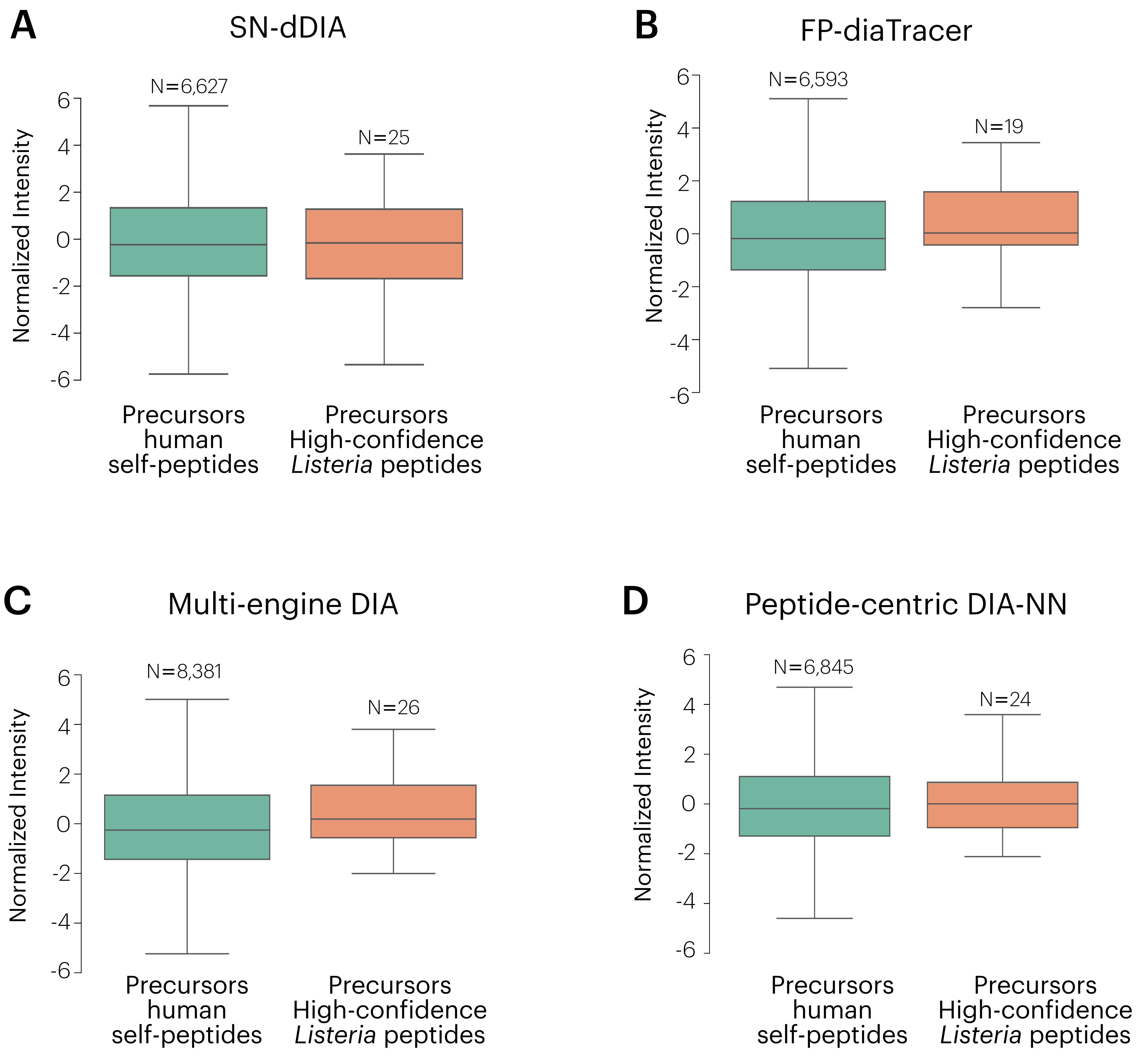

**Supplemental Figure S7. *Listeria* peptide precursor show similar abundance as human self-peptides.** Normalized intensity of human self-peptide and *Listeria* peptide precursors for each DIA analysis workflow: SN-dDIA (**A**), FP-diaTracer (**B**), multi-engine DIA (**C**) and peptide-centric DIA-NN workflows (**D**).

**Supplemental Table 1. DIA method generated by the py_diAID package^7^ after manual adjustment, using an in-house JY immunopeptidomics quality control sample.**

|  |  | **1/K_0_ (Vs/cm^2^)** | | **Mass (*m/z*)** | |
| --- | --- | --- | --- | --- | --- |
| **MS Type** | **Cycle Id** | **Begin** | **End** | **Start** | **End** |
| MS1 | 0 | - | - | - | - |
| PASEF | 1 | 1.1 | 1.65 | 826.42 | 857.41 |
| PASEF | 1 | 0.87 | 1.1 | 578.78 | 595.76 |
| PASEF | 1 | 0.75 | 0.87 | 365.22 | 464.26 |
| PASEF | 2 | 1.13 | 1.65 | 857.41 | 888.45 |
| PASEF | 2 | 0.92 | 1.13 | 595.76 | 614.79 |
| PASEF | 2 | 0.75 | 0.92 | 464.26 | 492.74 |
| PASEF | 3 | 1.14 | 1.65 | 888.45 | 916.51 |
| PASEF | 3 | 0.93 | 1.14 | 614.79 | 638.42 |
| PASEF | 3 | 0.75 | 0.93 | 492.74 | 510.73 |
| PASEF | 4 | 1.16 | 1.65 | 916.51 | 945.5 |
| PASEF | 4 | 0.95 | 1.16 | 638.42 | 667.69 |
| PASEF | 4 | 0.75 | 0.95 | 510.73 | 524.79 |
| PASEF | 5 | 1.17 | 1.65 | 945.5 | 976.34 |
| PASEF | 5 | 0.96 | 1.17 | 667.69 | 705.9 |
| PASEF | 5 | 0.75 | 0.96 | 524.79 | 538.23 |
| PASEF | 6 | 1.18 | 1.65 | 976.34 | 1011.55 |
| PASEF | 6 | 0.97 | 1.18 | 705.9 | 750.39 |
| PASEF | 6 | 0.75 | 0.97 | 538.23 | 550.99 |
| PASEF | 7 | 1.2 | 1.65 | 1011.55 | 1051.66 |
| PASEF | 7 | 0.98 | 1.2 | 750.39 | 791.43 |
| PASEF | 7 | 0.75 | 0.98 | 550.99 | 564.34 |
| PASEF | 8 | 1.26 | 1.65 | 1051.66 | 1240.66 |
| PASEF | 8 | 1.01 | 1.26 | 791.43 | 826.42 |
| PASEF | 8 | 0.75 | 1.01 | 564.34 | 578.78 |
