## Supplementary material for "Data-independent immunopeptidomics discovery of low-abundant bacterial epitopes": Data S6

**Supplemental Data S6. Chromatograms extracted from DIA-NN Viewer or Spectronaut 19 for the 37 identified high-quality Listeria peptides.** Peptides were sorted alphabetically. Also see Supplemental Data S5 for the list of high-confident Listeria peptides.

AFIKIIQLRKNN/3+, UniProtKB A0A3Q0NB12 matching LMON_0202, actA, Actin-assembly inducing protein ActA.

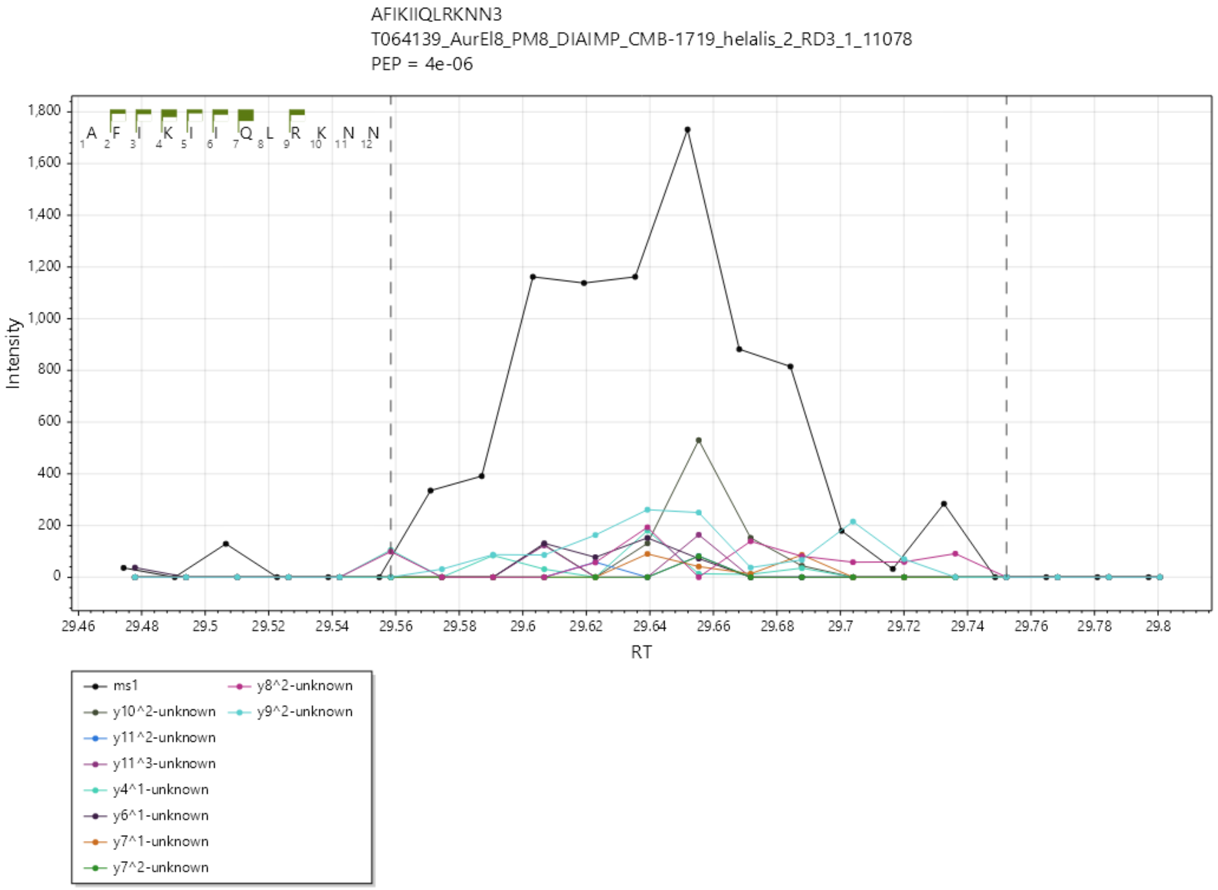

AKYFNQSVTDY/2+, UniProtKB A0A3Q0NAV2 matching LMON_0203, plcB, Phospholipase C.

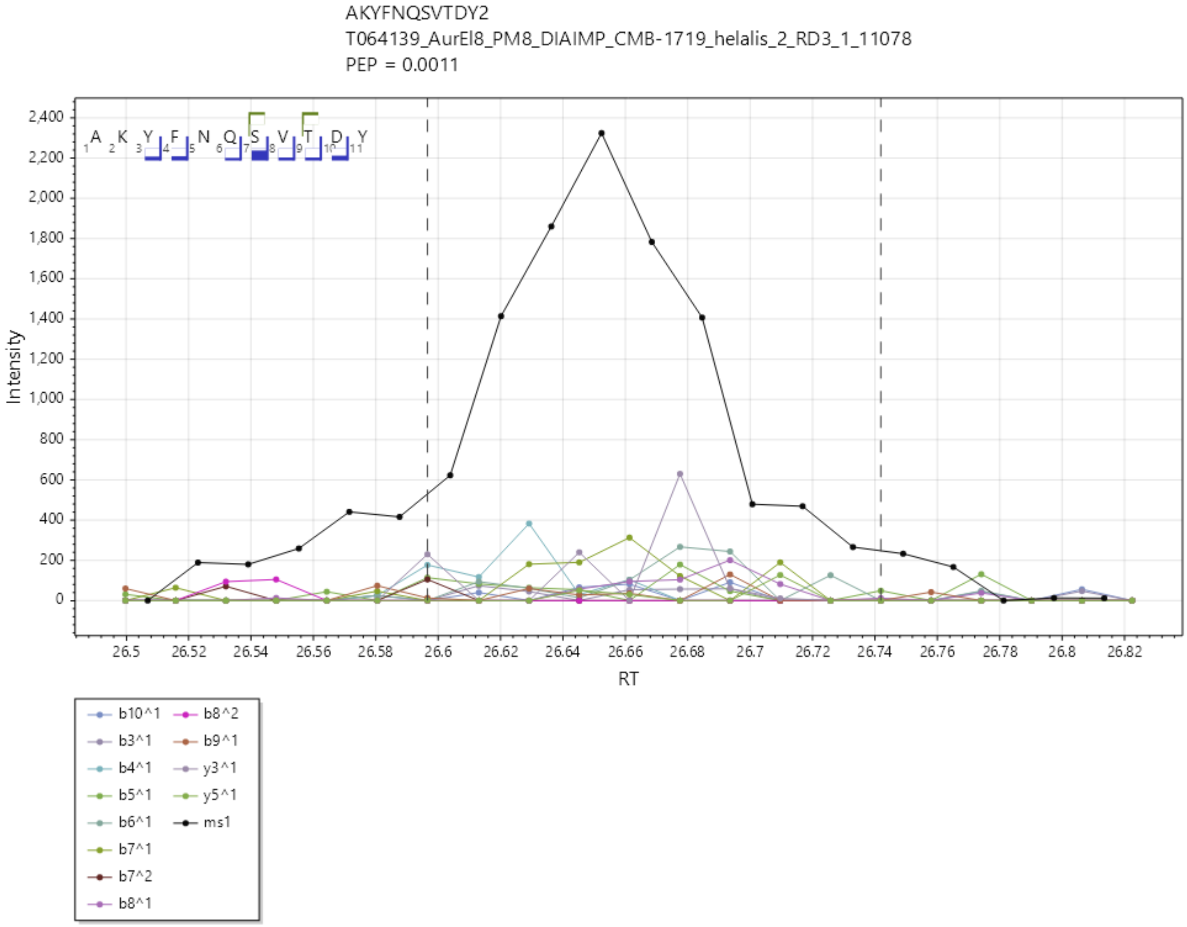

DLFDRAFVV/2+, UniProtKB A0A3Q0NEI0 matching LMON_1501, ftsI, Cell division protein FtsI [Peptidoglycan synthetase] / Transpeptidase, Penicillin binding protein transpeptidase domain.

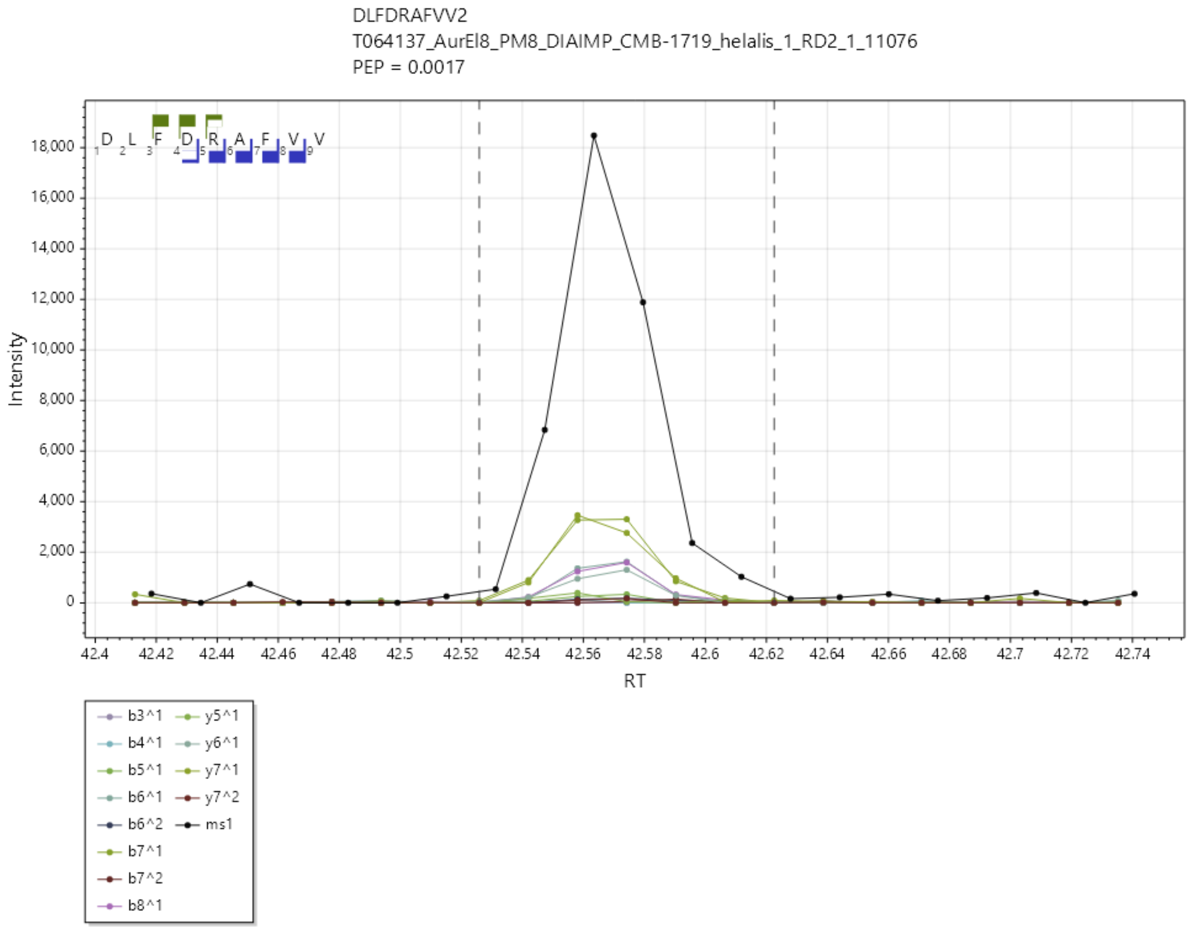

EIAIPTLFV/1+, UniProtKB A0A3Q0ND25 matching LMON_0957, Cell surface hydrolase, membrane-bound.

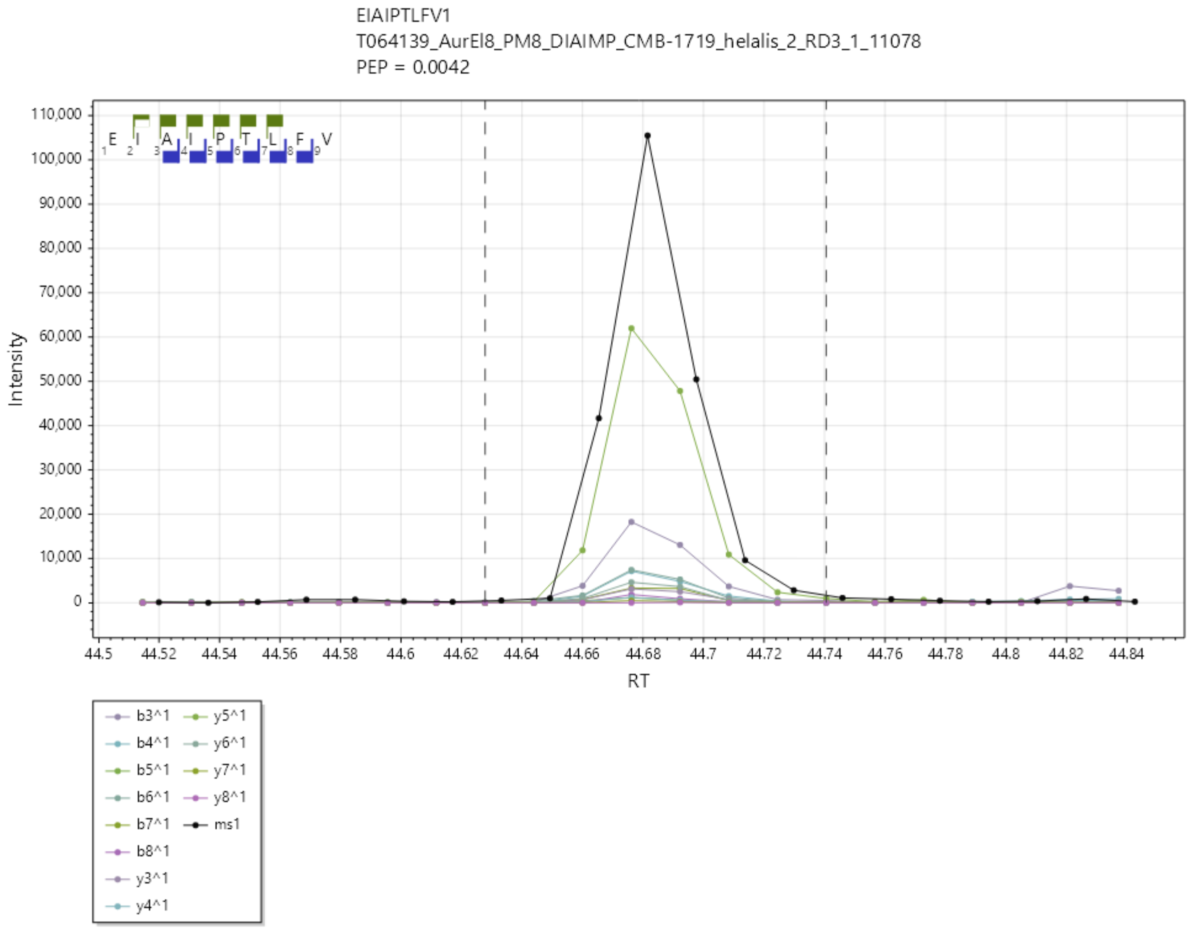

EIFDVKVAKV/2+, UniProtKB A0A3Q0NHR6 matching LMON_2651, rplW, Large ribosomal subunit protein uL23.

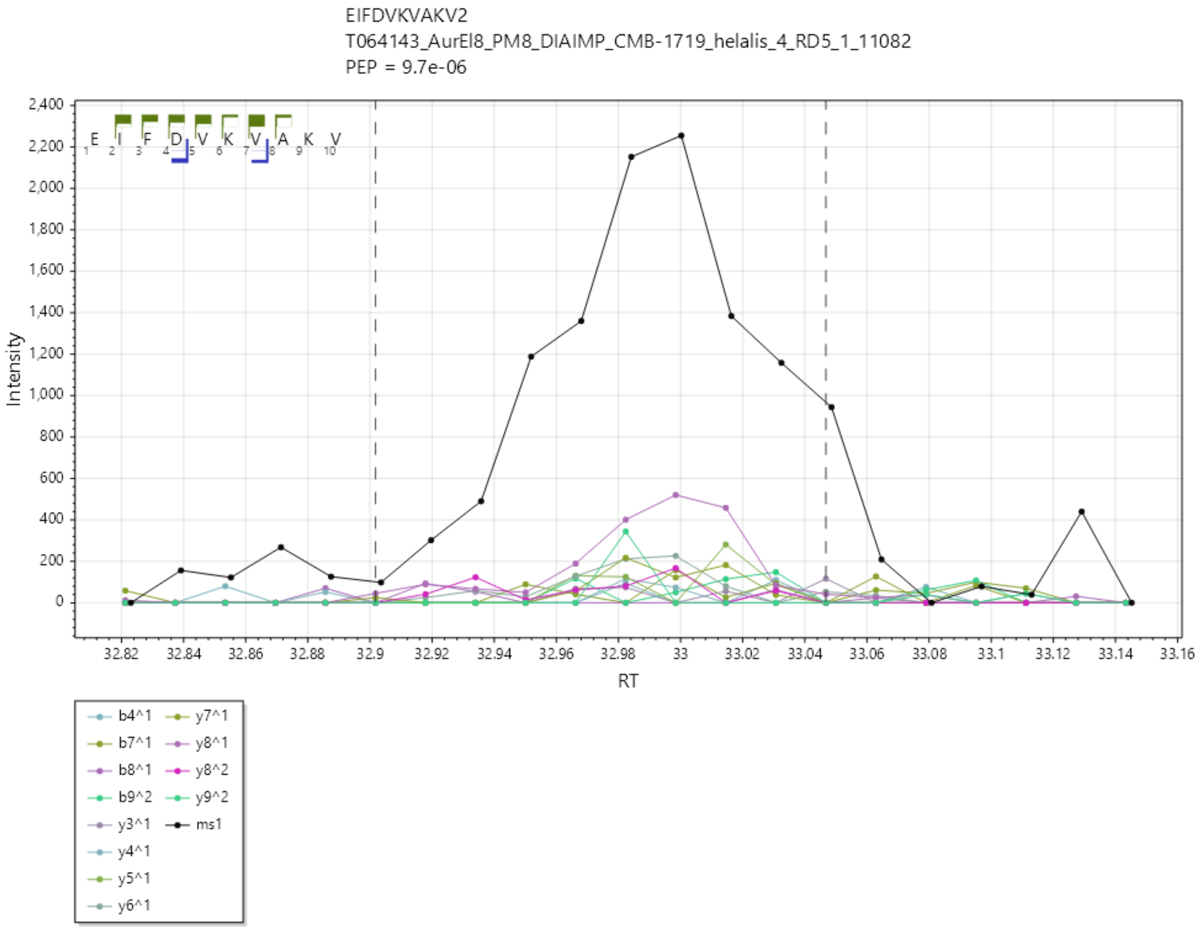

ESFSNVVTI/2+, UniProtKB A0A3Q0NAS1, LMON_0179, N-Acetyl-D-glucosamine ABC transport system, sugar-binding protein.

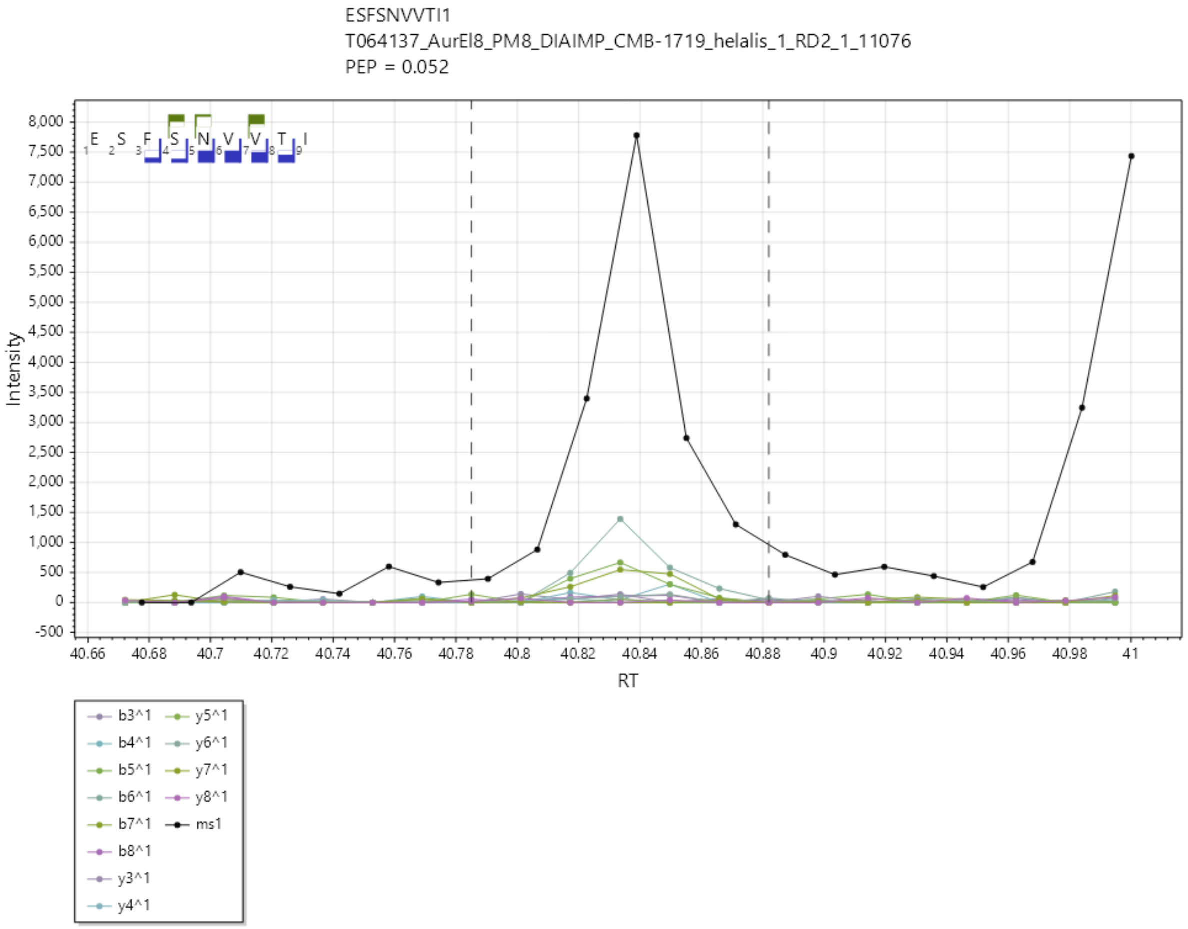

ETFFPQKESYV/2+, UniProtKB A0A3Q0NAQ5, LMON_0149, OppA, Oligopeptide ABC transporter, periplasmic oligopeptide-binding protein OppA (TC 3.A.1.5.1).

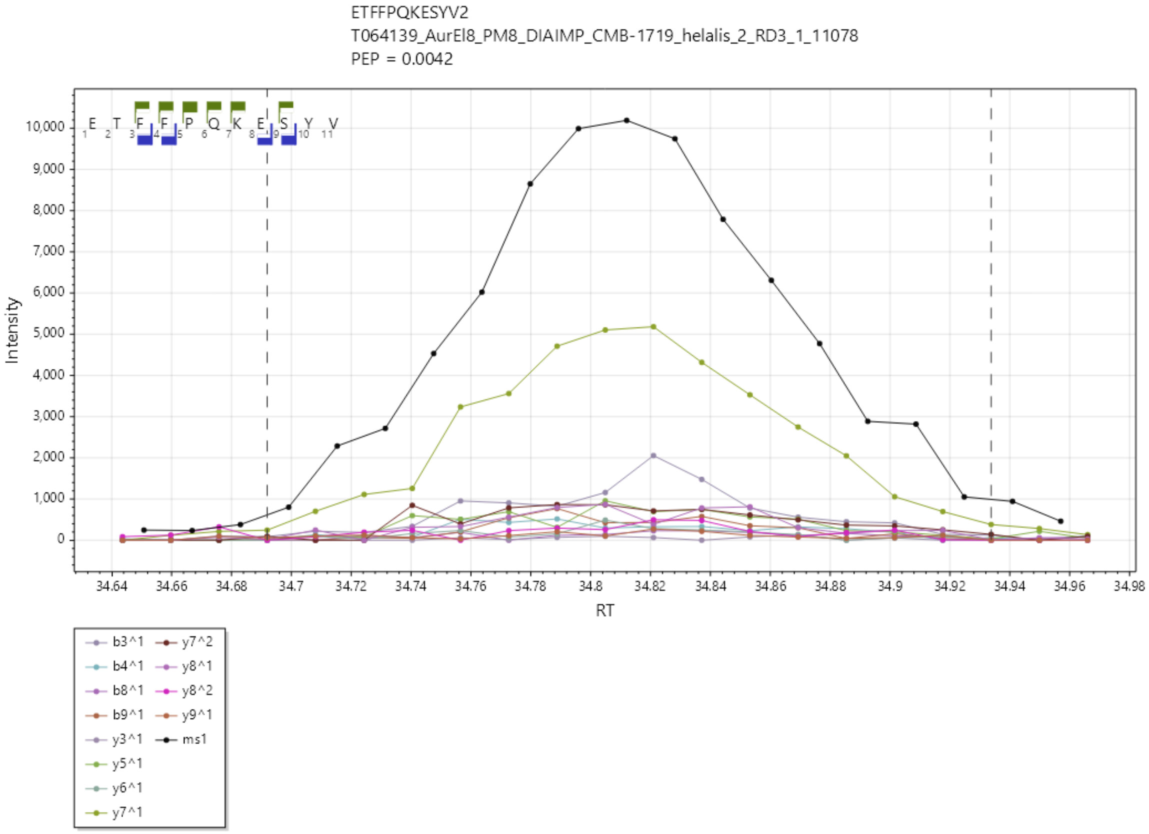

ETINKLIRV/2+, UniProtKB A0A3Q0NEI5 matching LMON_1517, rpoD, RNA polymerase sigma factor SigA.

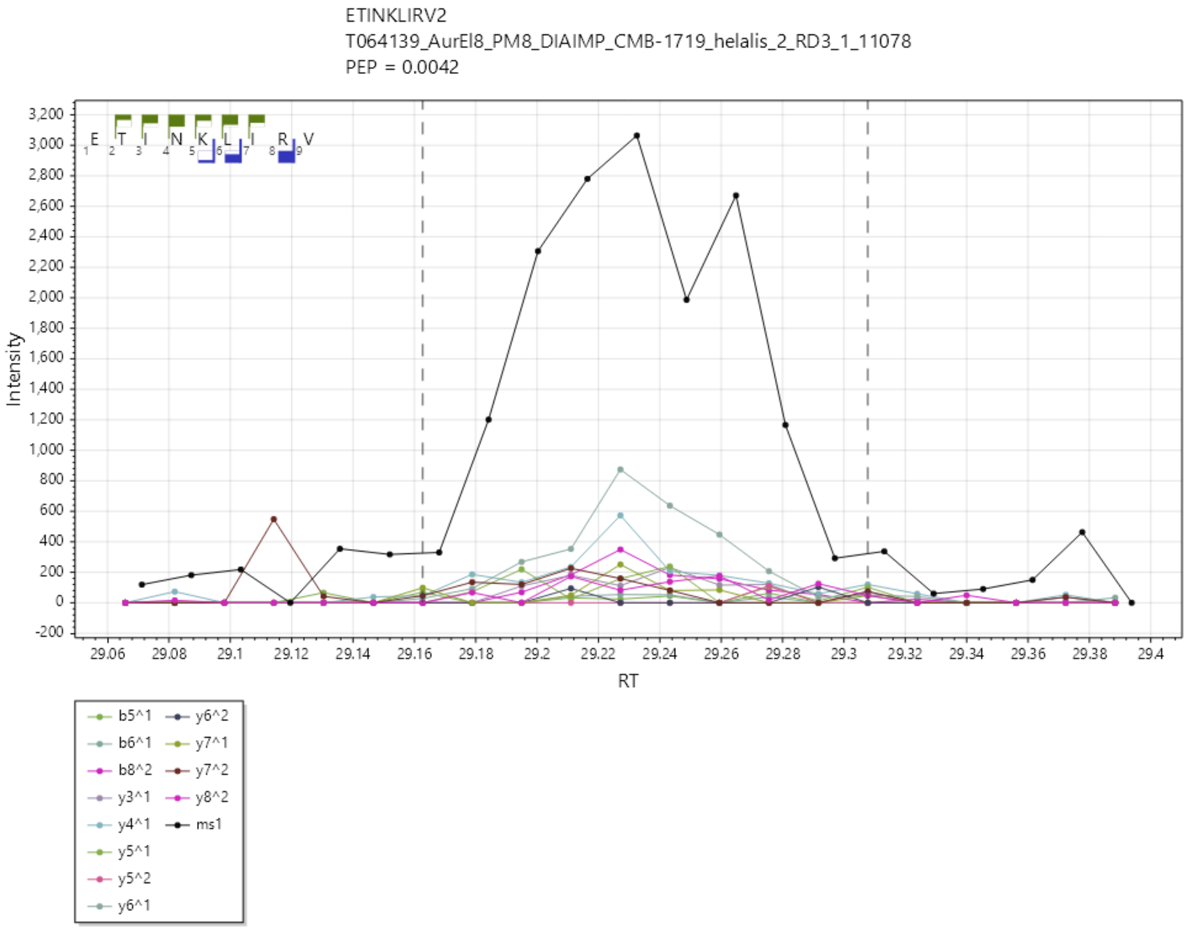

EVASWIVQV/1+, UniProtKB A0A3Q0NAW3 matching LMON_0201, mpl, Neutral metalloproteinase.

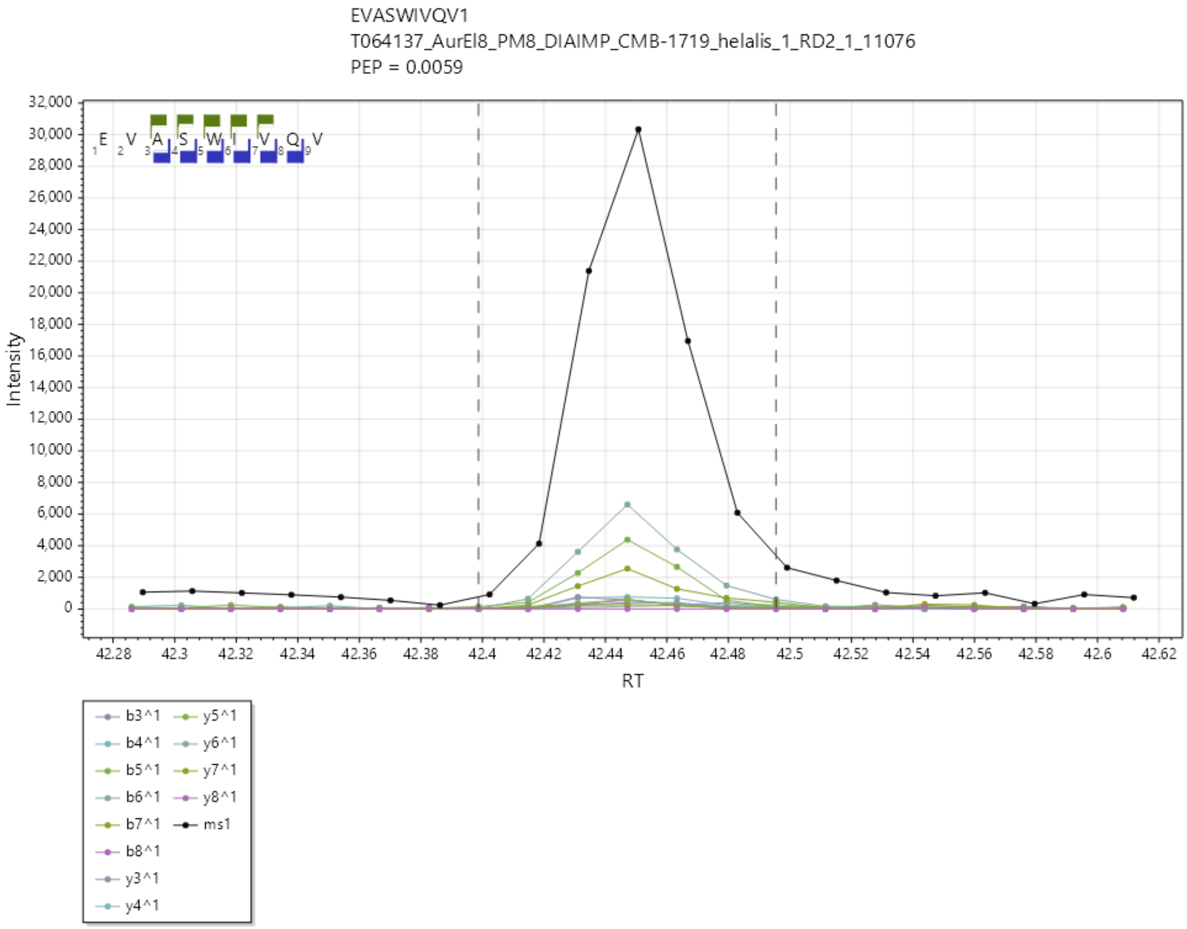

EVERPSLGV/2+, UniProtKB A0A3Q0NB40 matching LMON_0294, DegP/HtrA, Serine protease, DegP/HtrA, do-like.

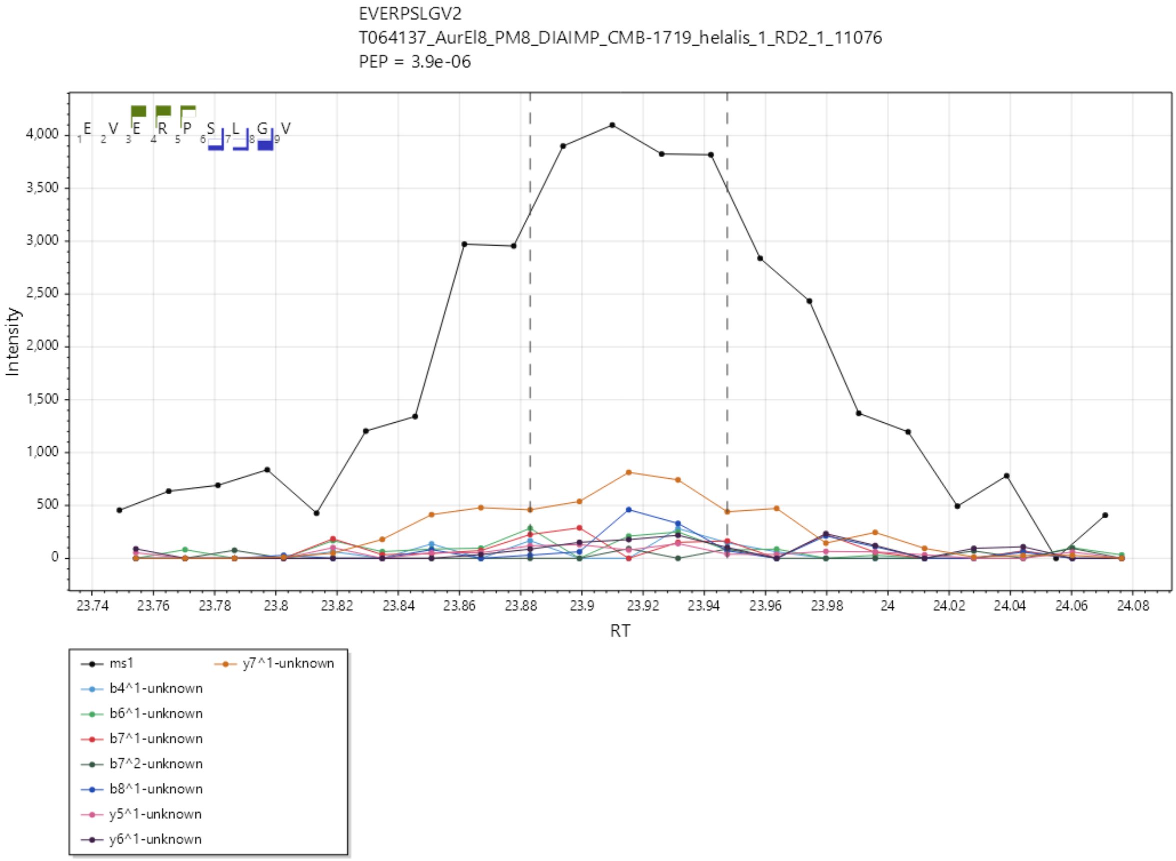

EVFDKYFEL/2+, UniProtKB A0A3Q0NEI0 matching LMON_1501, ftsI, Cell division protein FtsI [Peptidoglycan synthetase] / Transpeptidase, Penicillin binding protein transpeptidase domain.

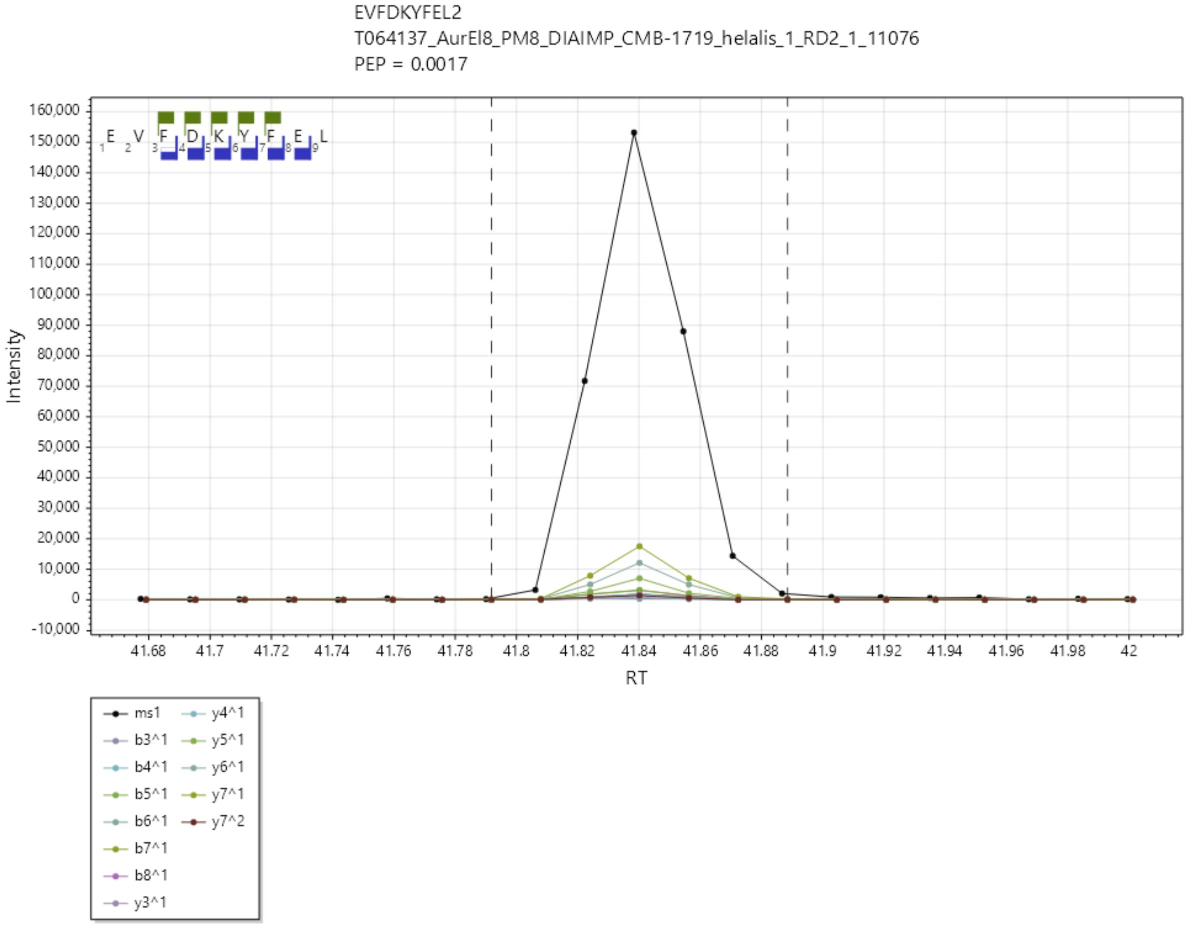

EVFDKYFELK/2+, UniProtKB A0A3Q0NEI0, matching LMON_1501, ftsI, Cell division protein FtsI [Peptidoglycan synthetase] / Transpeptidase, Penicillin binding protein transpeptidase domain.

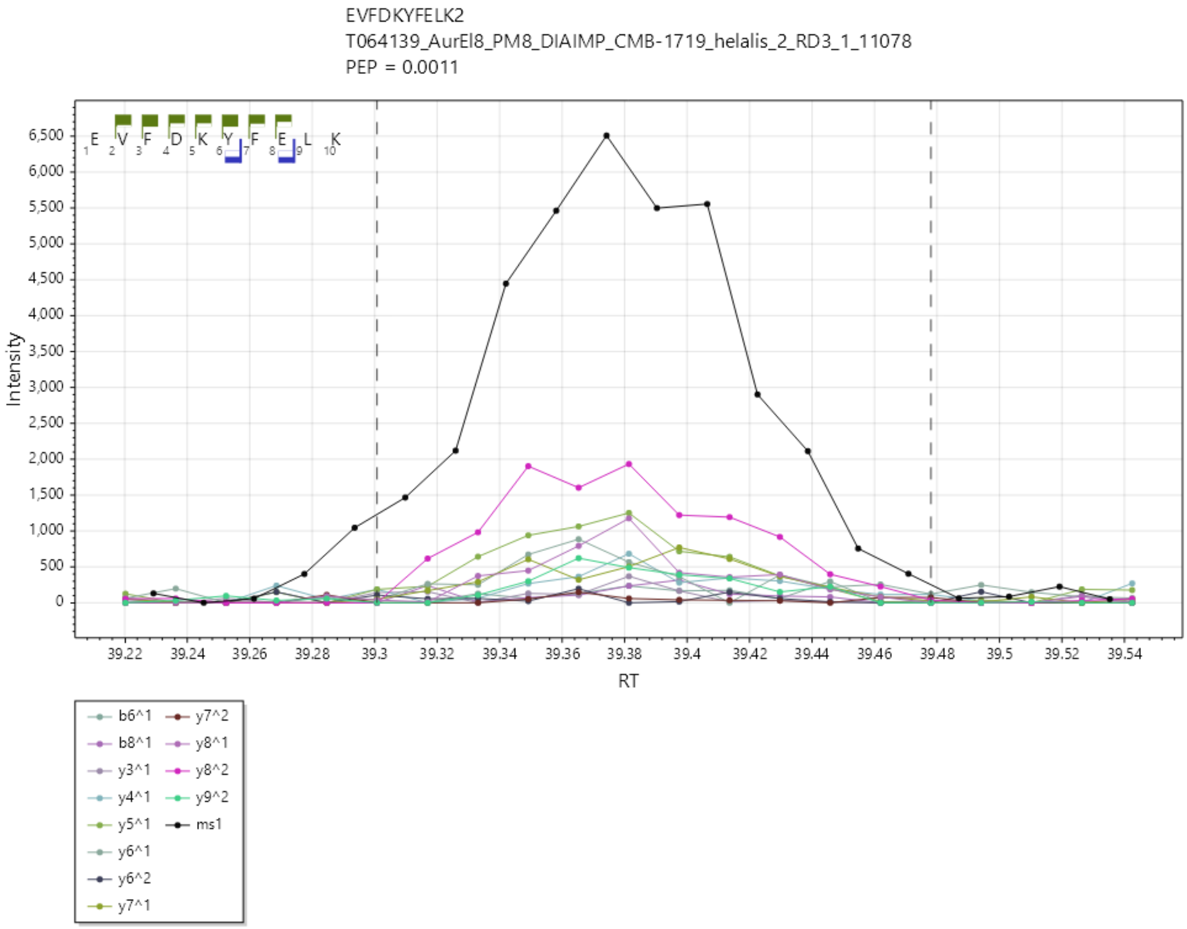

EVIGTAILI/1+, UniProtKB A0A3Q0NF08 matching LMON_1605, LMON_1605, Glycerol uptake facilitator protein

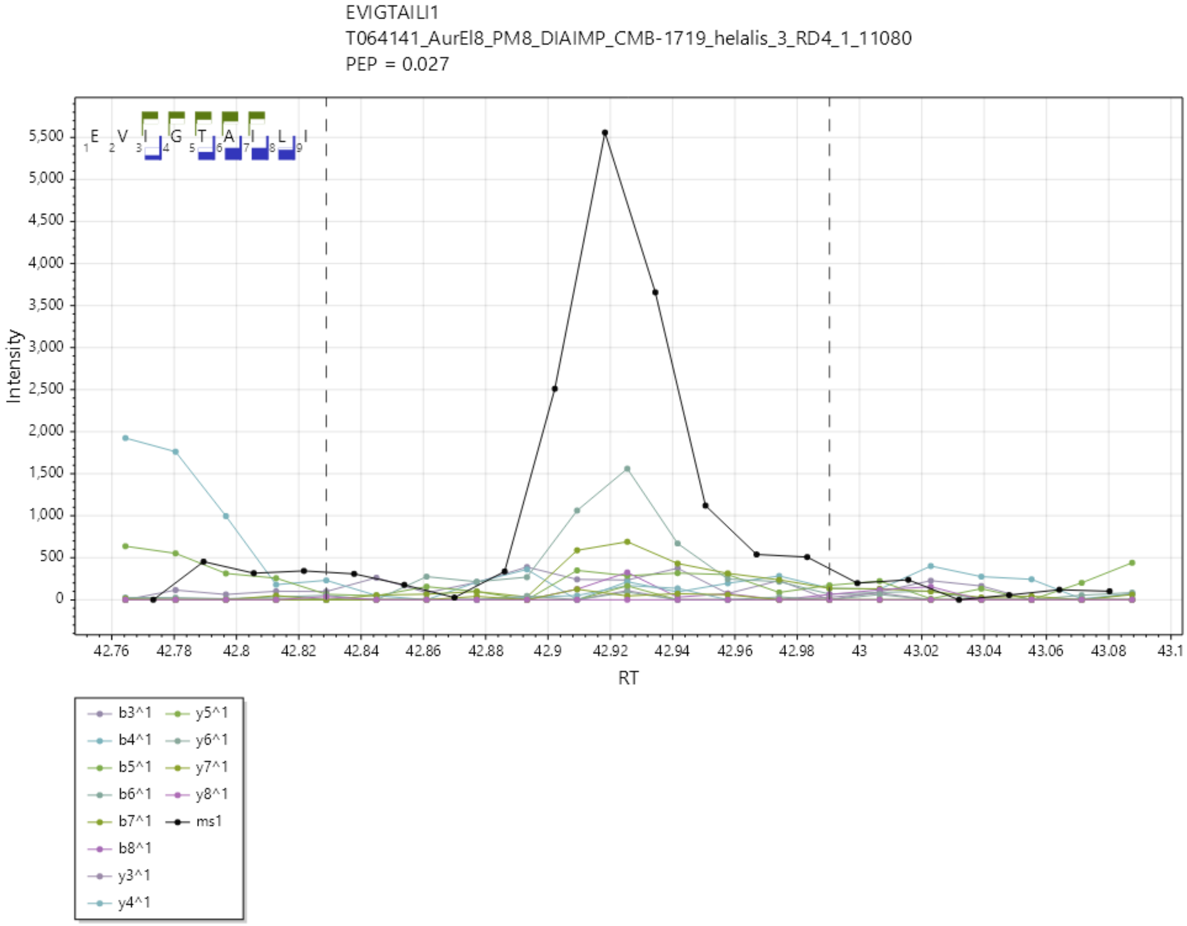

EVIGVIAGL/1+, UniProtKB A0A3Q0NEE4 matching LMON_1421, Alkaline shock protein.

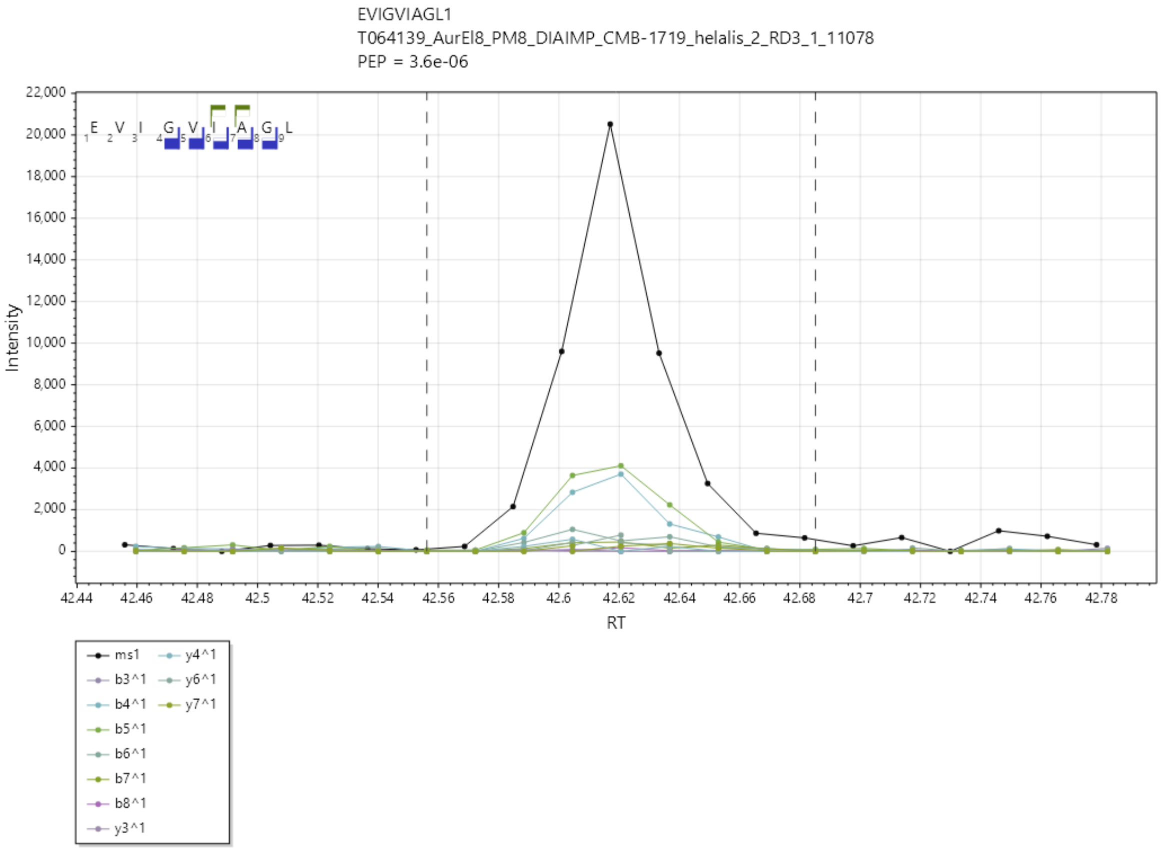

EVPEILLEV/1+, UniProtKB A0A3Q0NHG8 matching LMON_2576, argS, Arginine--tRNA ligase.

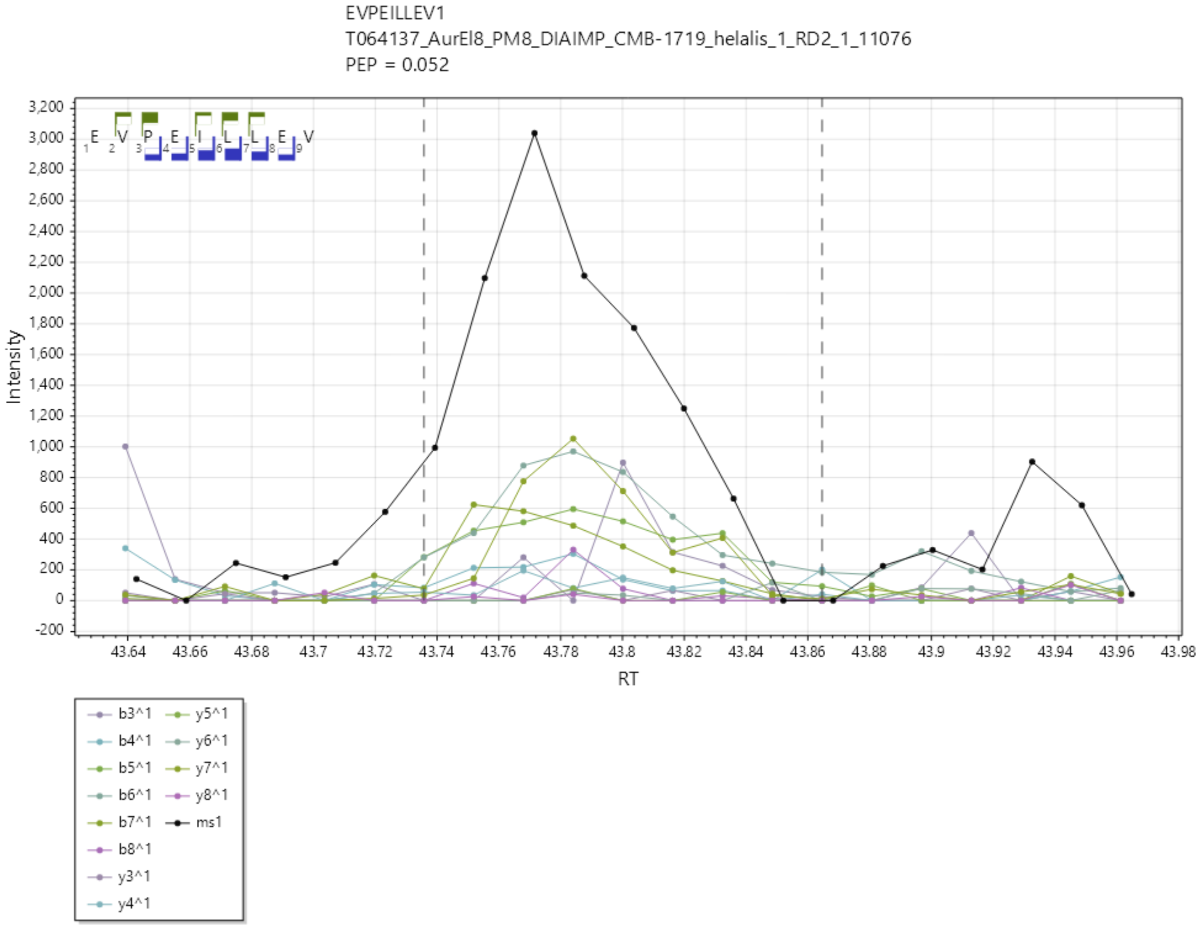

FAAPTIASA/1+ UniProtKB A0A3Q0NBW8 matching LMON_0582, iap, Probable endopeptidase p60.

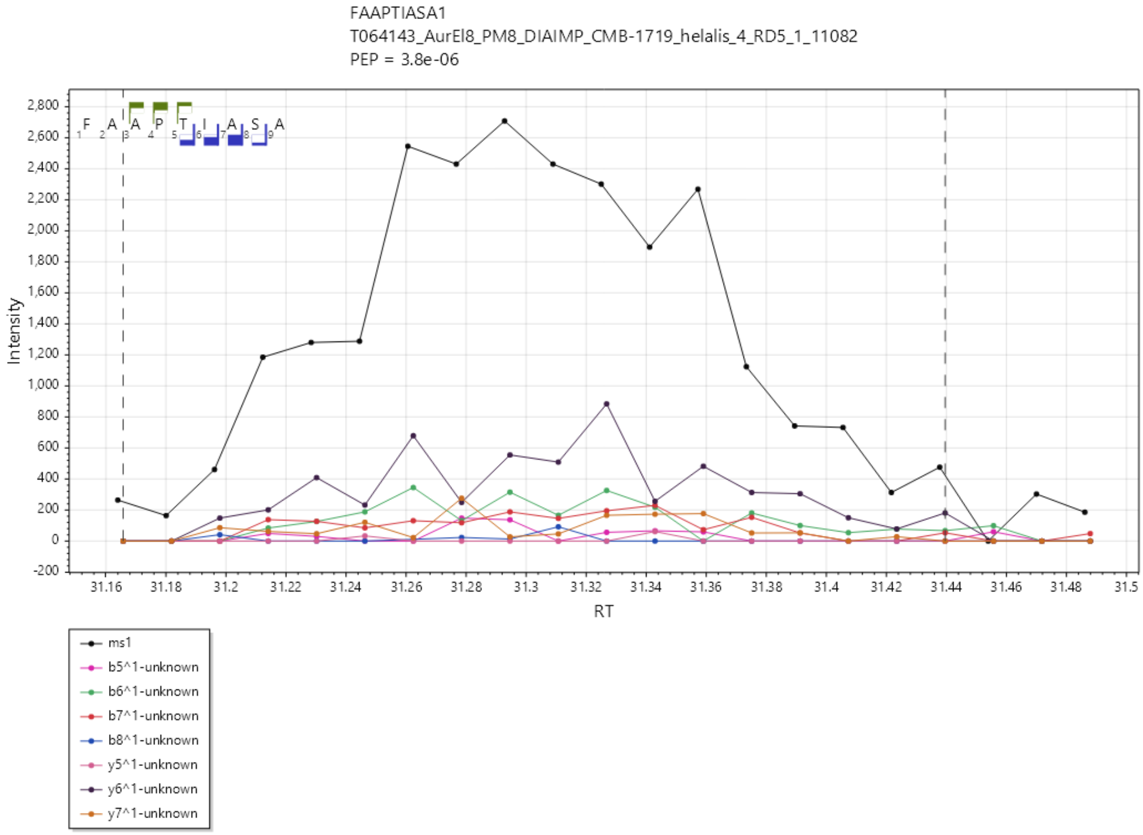

FDKYFEL/1+, UniProtKB A0A3Q0NEI0 matching LMON_1501, ftsI, Cell division protein FtsI [Peptidoglycan synthetase] / Transpeptidase, Penicillin binding protein transpeptidase domain.

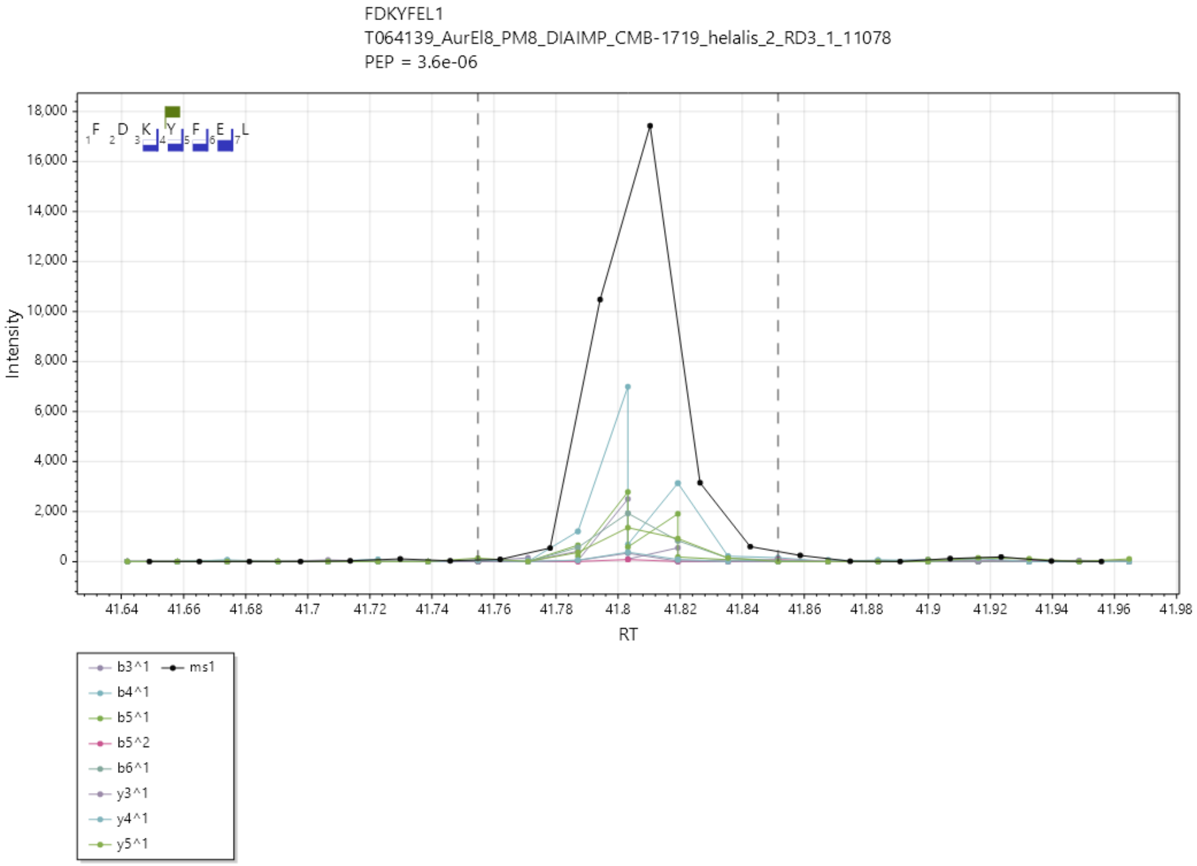

FVFDNKPVKV/2+, UniProtKB A0A3Q0NAM8 matching LMON_0134, OppA*, Oligopeptide ABC transporter, periplasmic oligopeptide-binding protein OppA (TC 3.A.1.5.1).

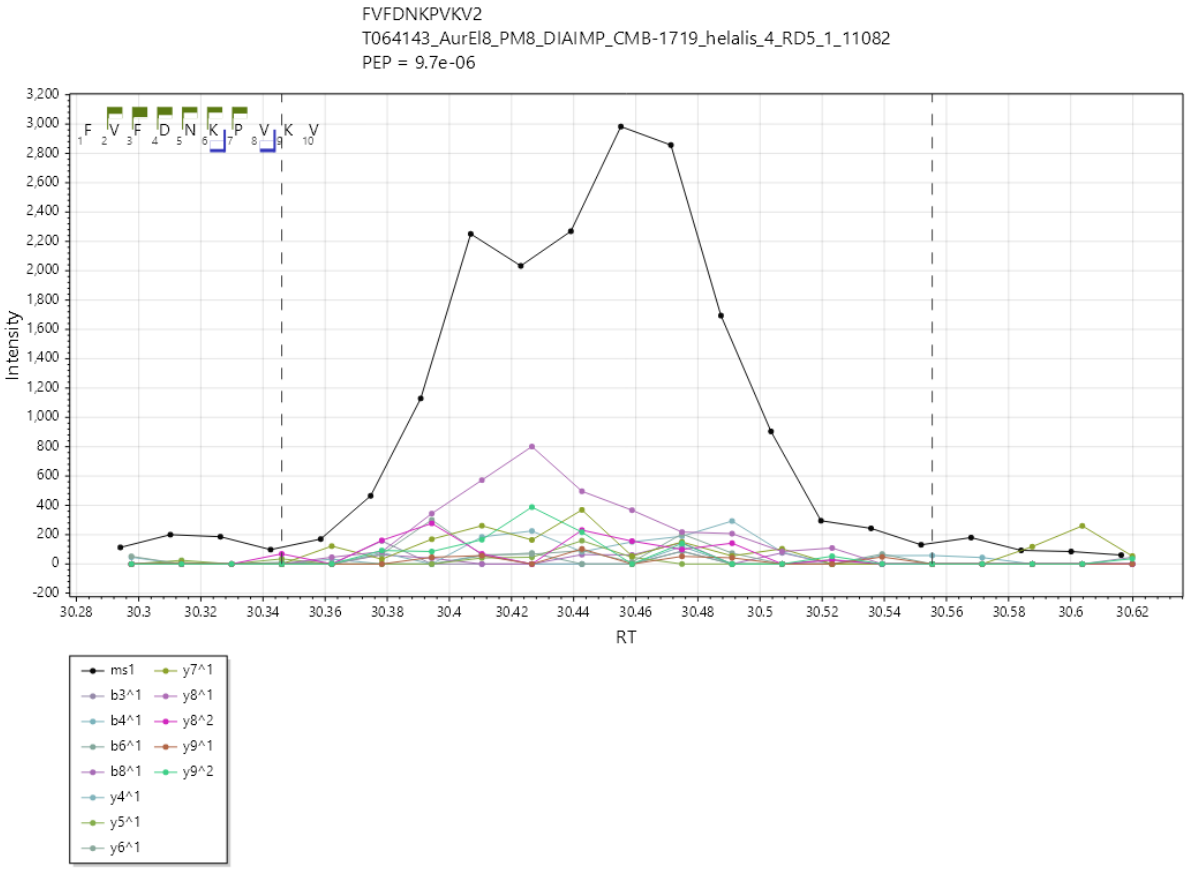

HLPEFTNEV/2+, UniProtKB A0A3Q0NBH6 matching LMON_0442, inlB, Internalin B (GW modules).

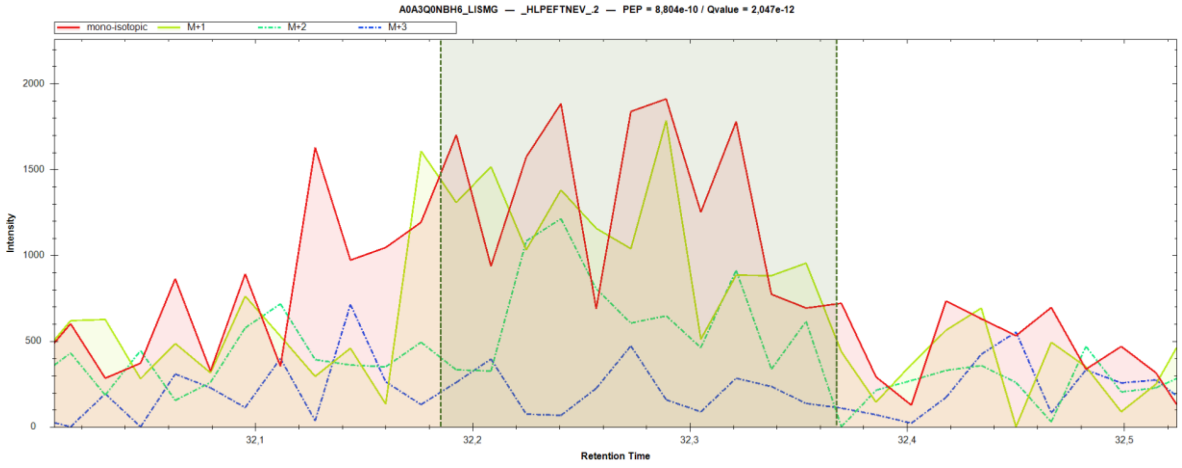


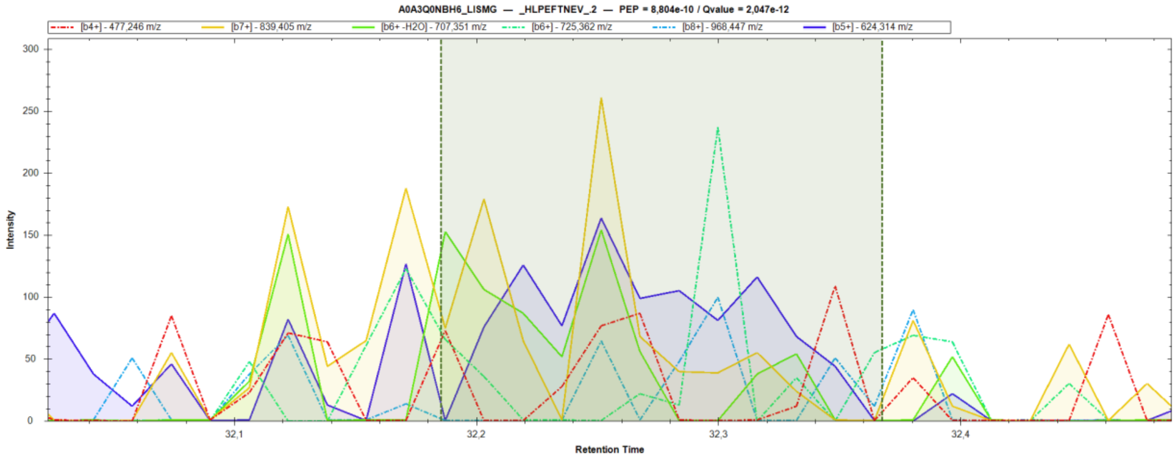

HTYAGEVKAIRGV/3+, UniProtKB A0A3Q0NGQ2 matching LMON_2269, OppD, Oligopeptide transport ATP-binding protein OppD (TC 3.A.1.5.1).

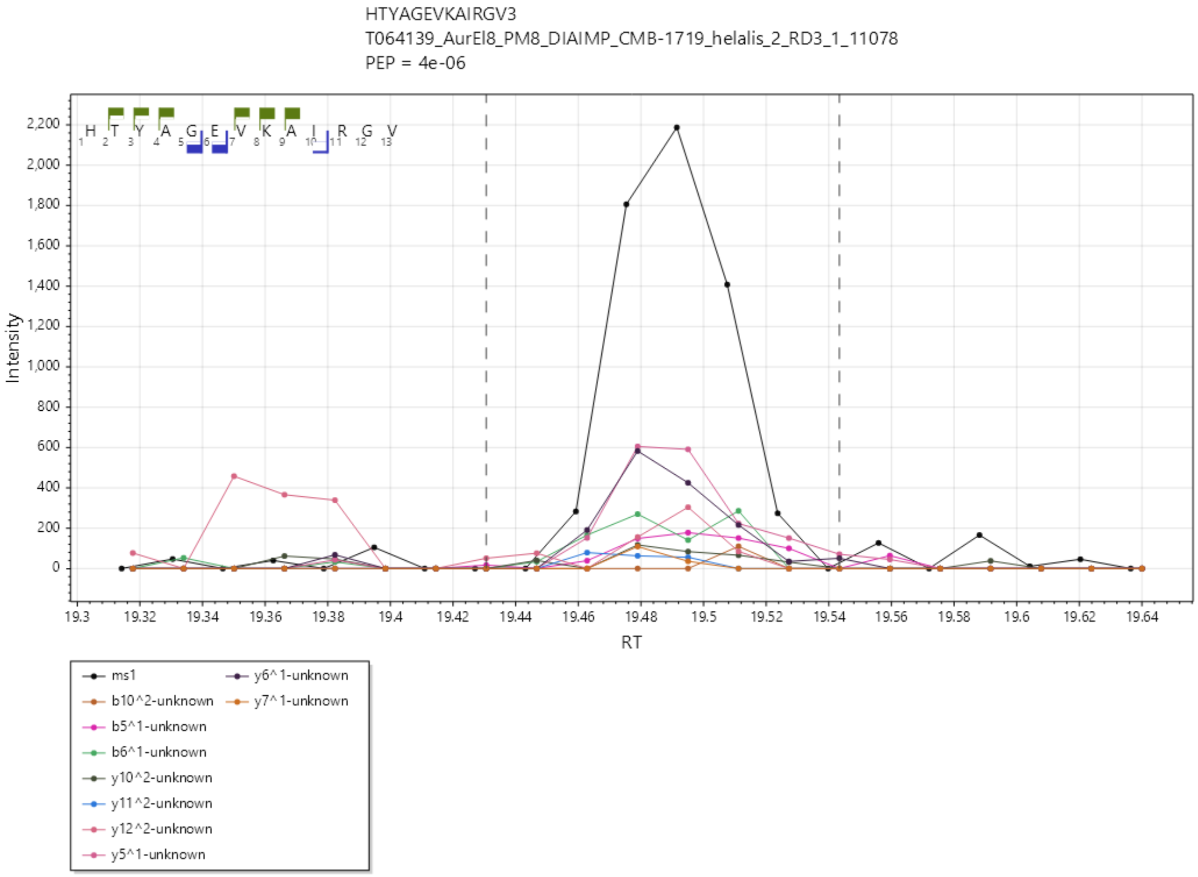

HVNIGTIGHV/2+, UniProtKB A0A3Q0NHS2 matching LMON_2676, tuf, Elongation factor Tu.

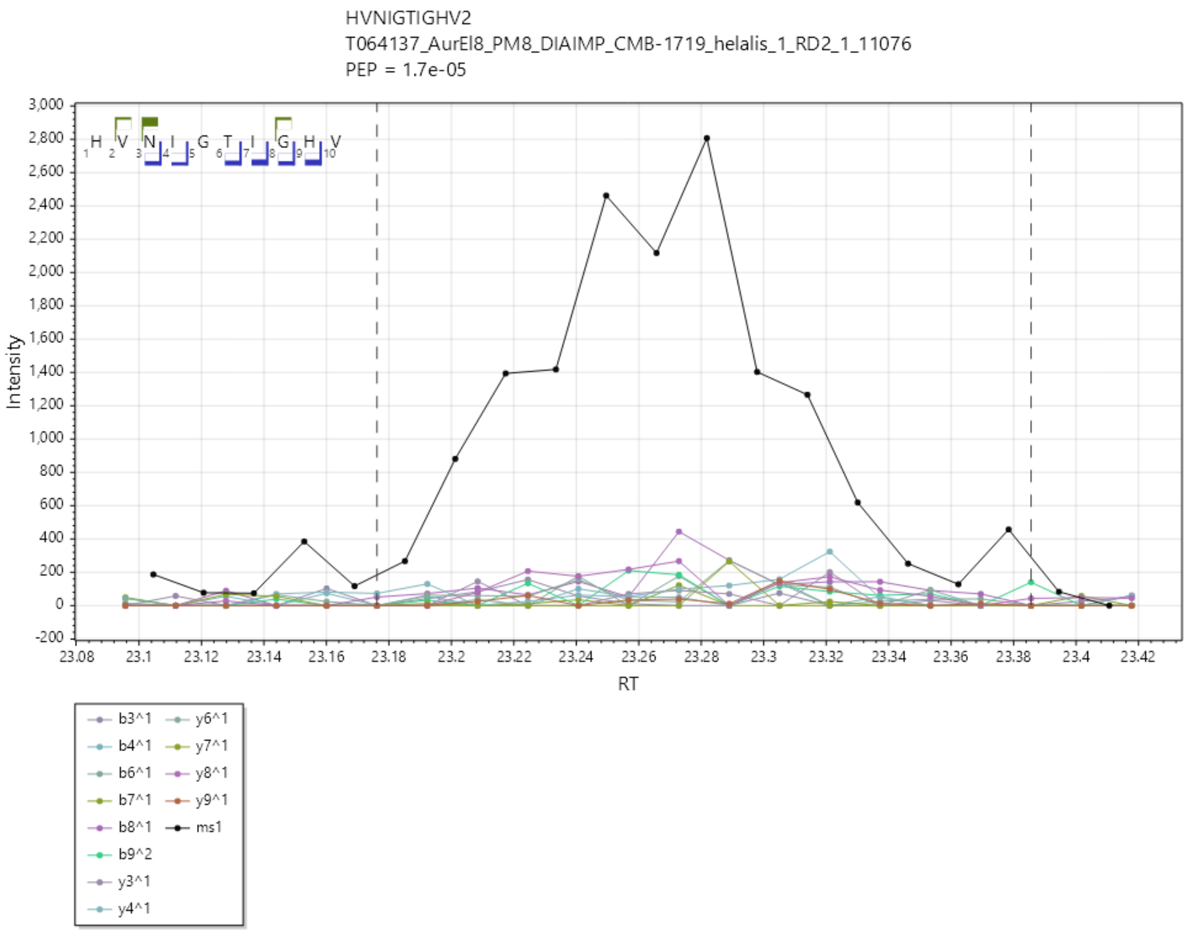

ILADIVKNV/2+, UniProtKB A0A3Q0NGJ0 matching LMON_1915, SitA, Manganese ABC transporter, periplasmic-binding protein SitA.

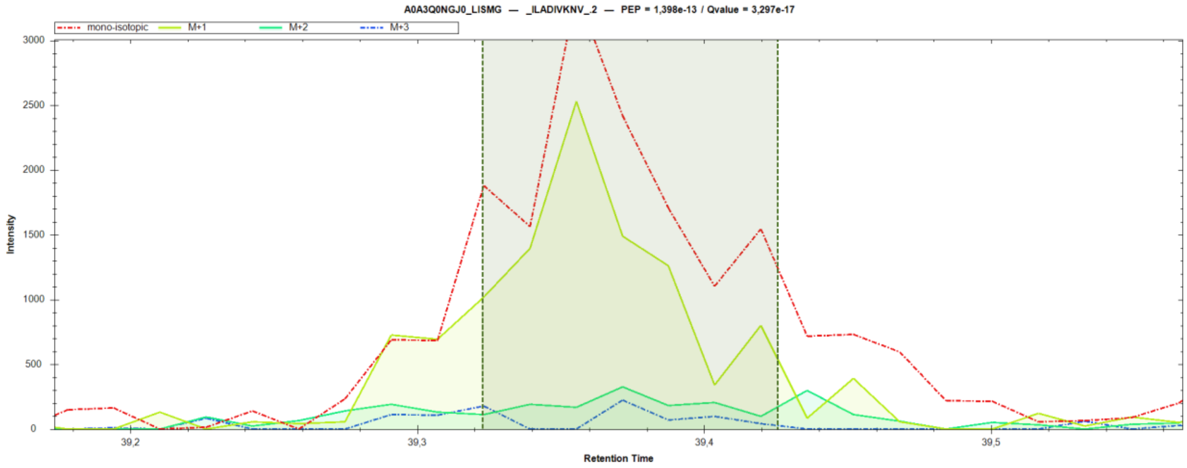


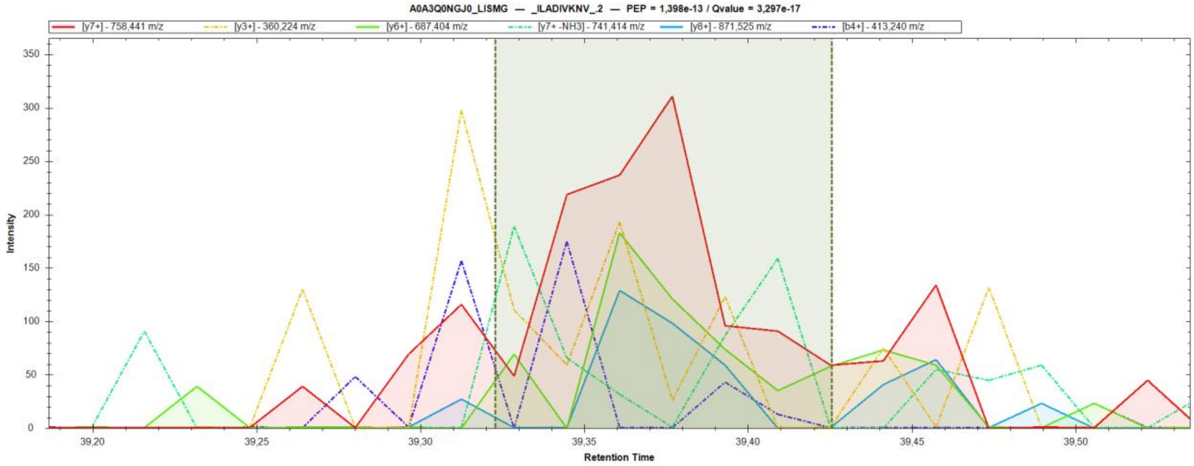

IAYGEAILL/1+, UniProtKB A0A3Q0NFF8 matching LMON_1812, BceB, Bacitracin export permease protein BceB.

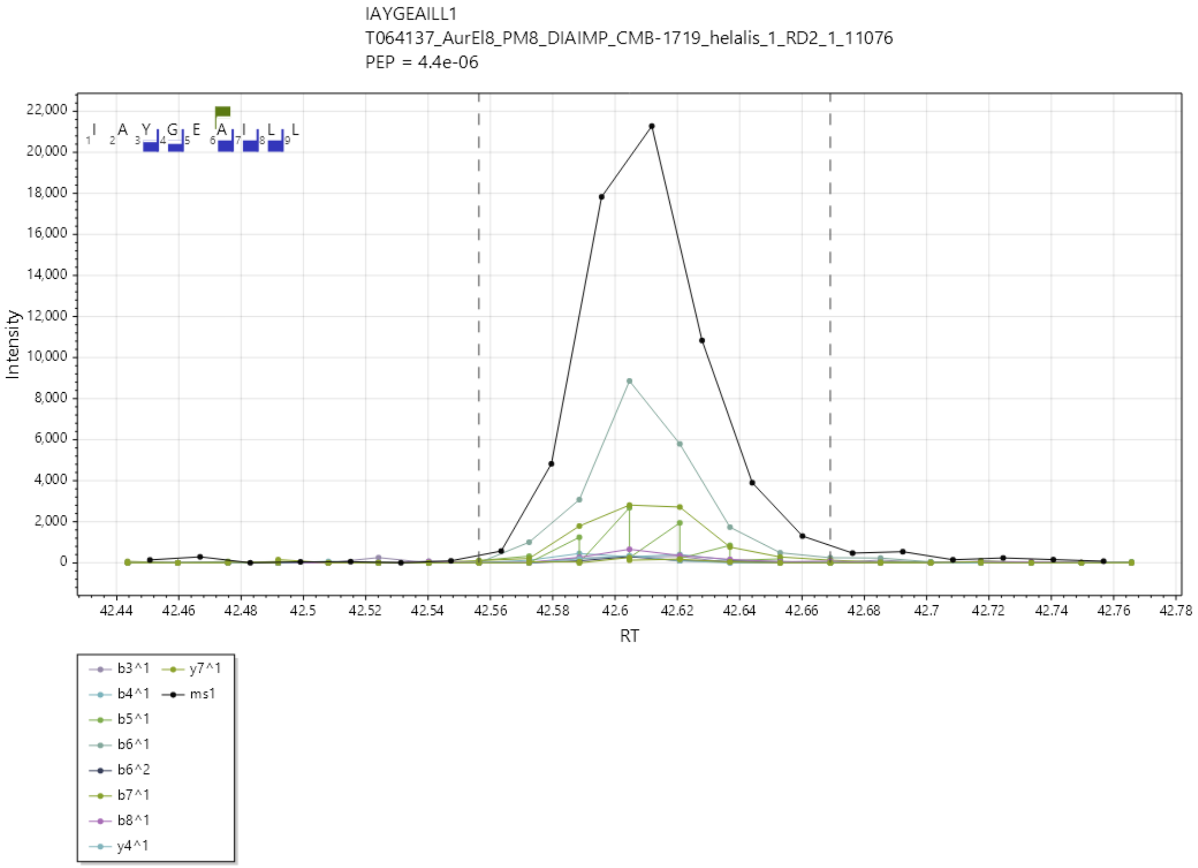

KLNDLISRY/2+, UniProtKB A0A3Q0NHX2 matching LMON_2714, LMON_2714, Peptidoglycan hydrolase.

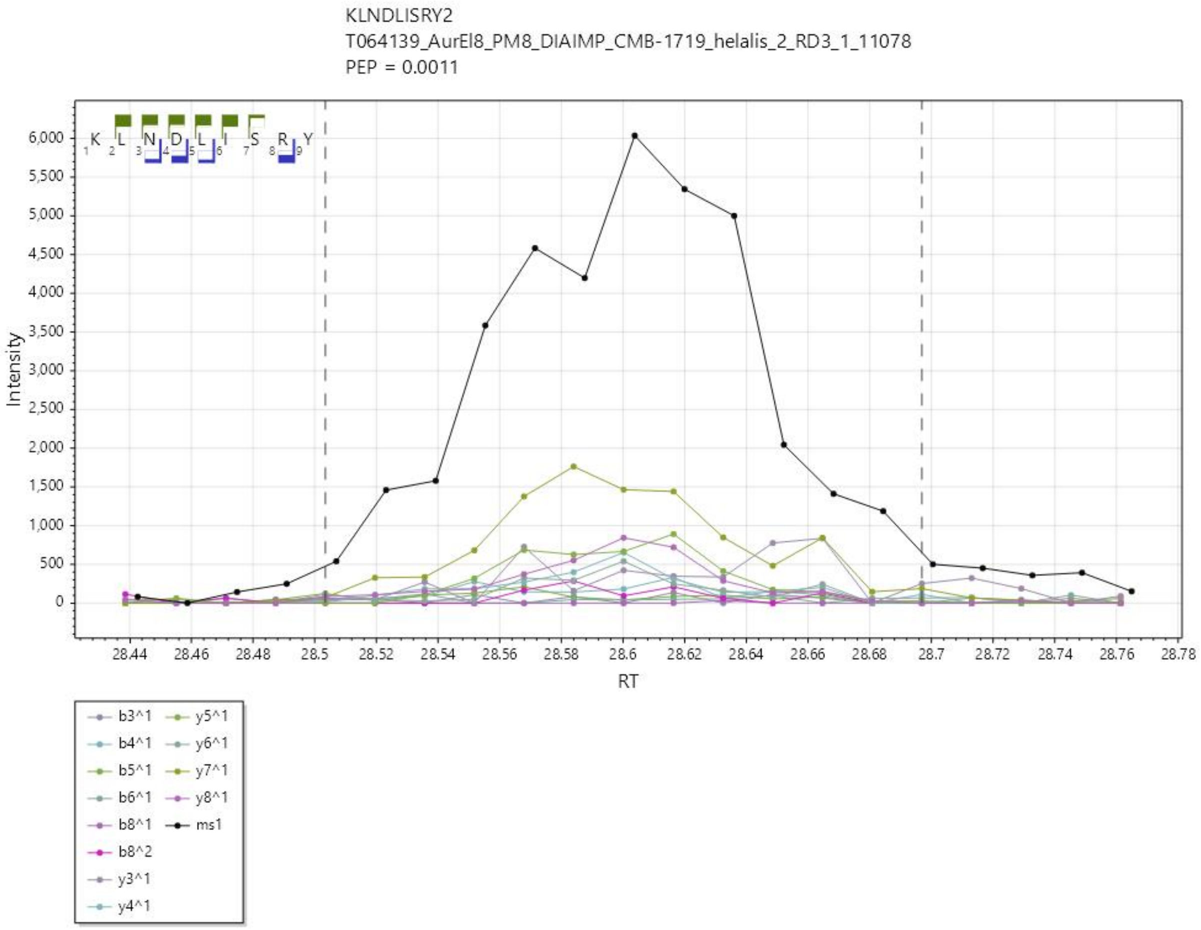

KMoxQEEVISF/2+, UniProtKB A0A3Q0NAV6 matching LMON_0200, hly, Thiol-activated cytolysin.

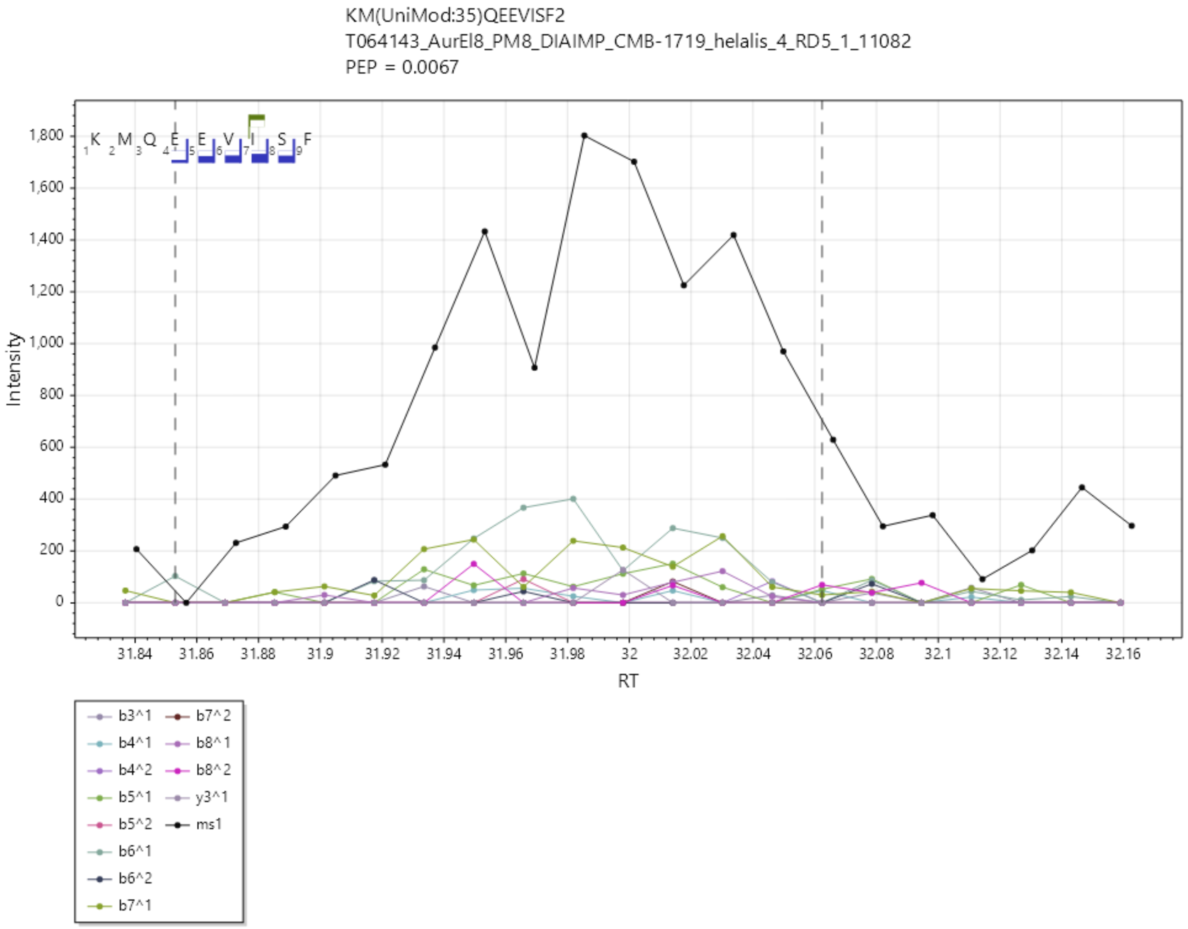

KVYDGILKV/2+, UniProtKB A0A3Q0NAY2 matching LMON_0221, cysK, Cysteine synthase.

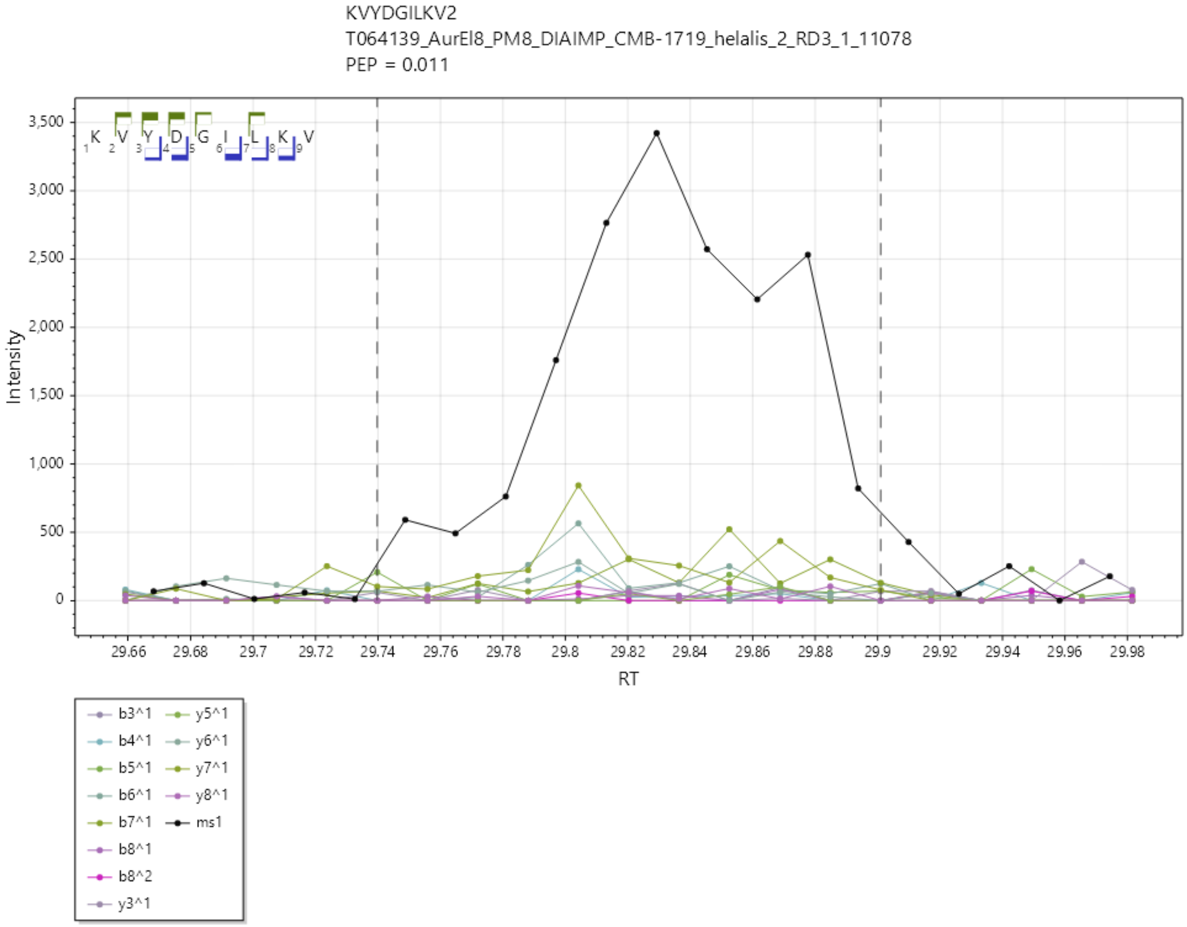

LEKTLGITV/2+, UniProtKB A0A3Q0NFM2 matching LMON_1883, LMON_1883, FIG001802: Putative alkaline-shock protein.

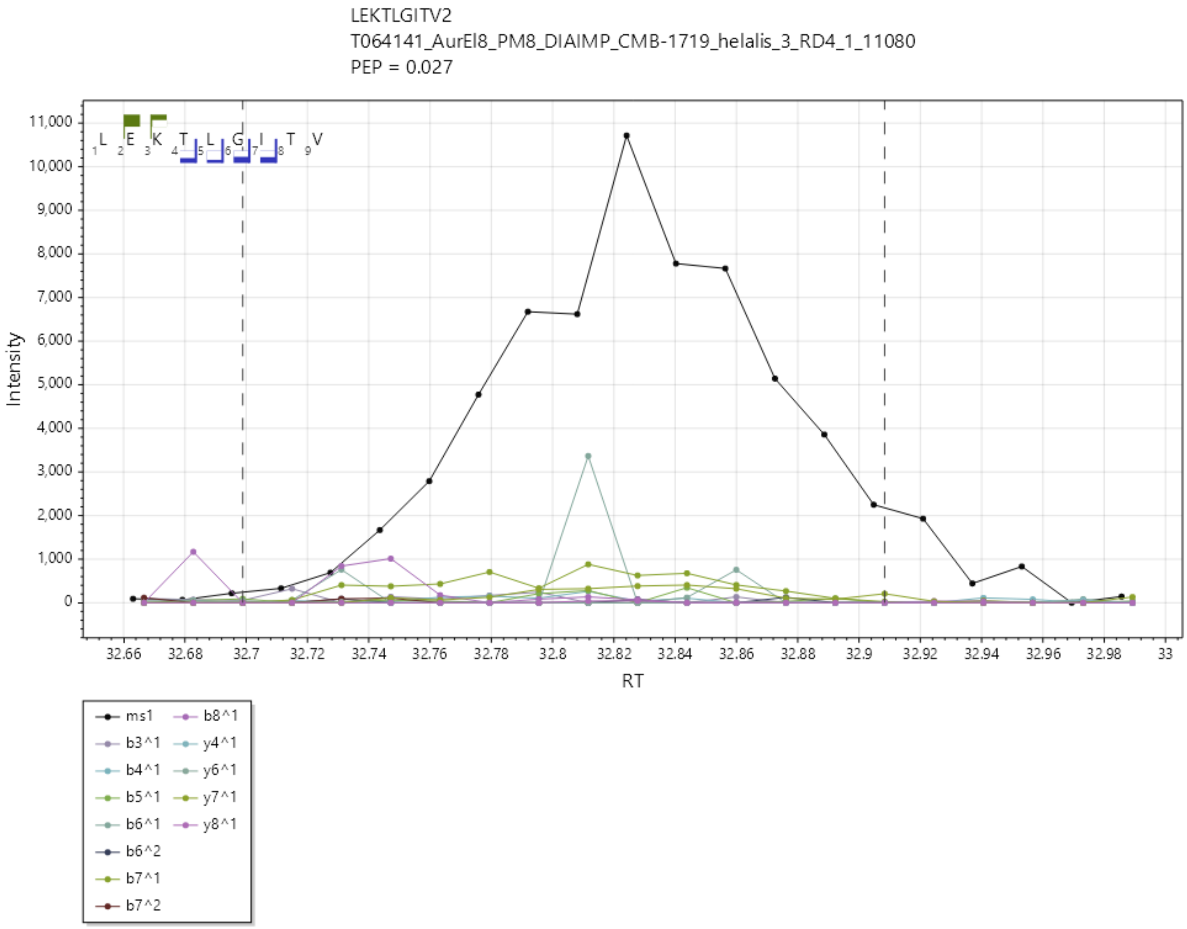

QVFEGLYTL/1+, UniProtKB A0A3Q0NAQ5 matching LMON_0149, OppA, Oligopeptide ABC transporter, periplasmic oligopeptide-binding protein OppA (TC 3.A.1.5.1).

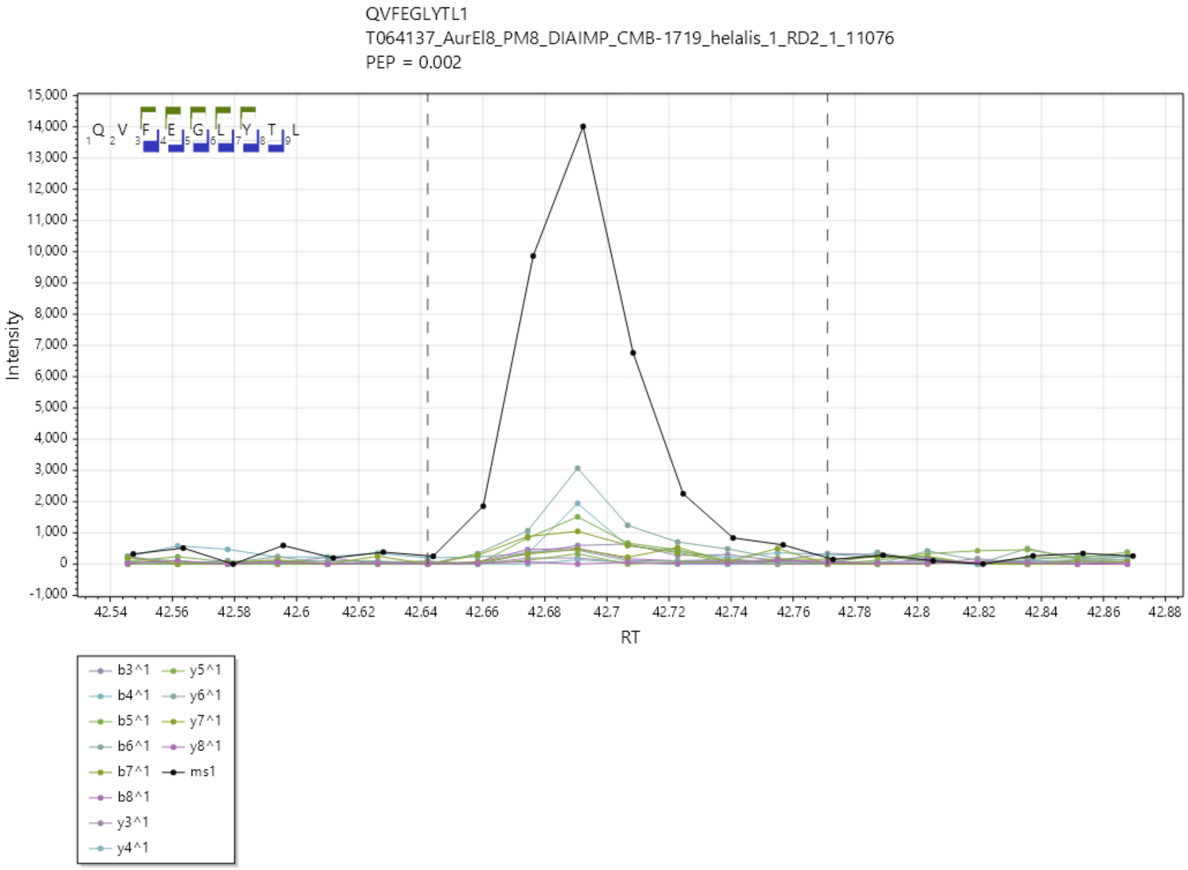

SINMoxPSLPV/1+, UniProtKB A0A3Q0NB12 matching LMON_0202, actA, Actin-assembly inducing protein ActA.

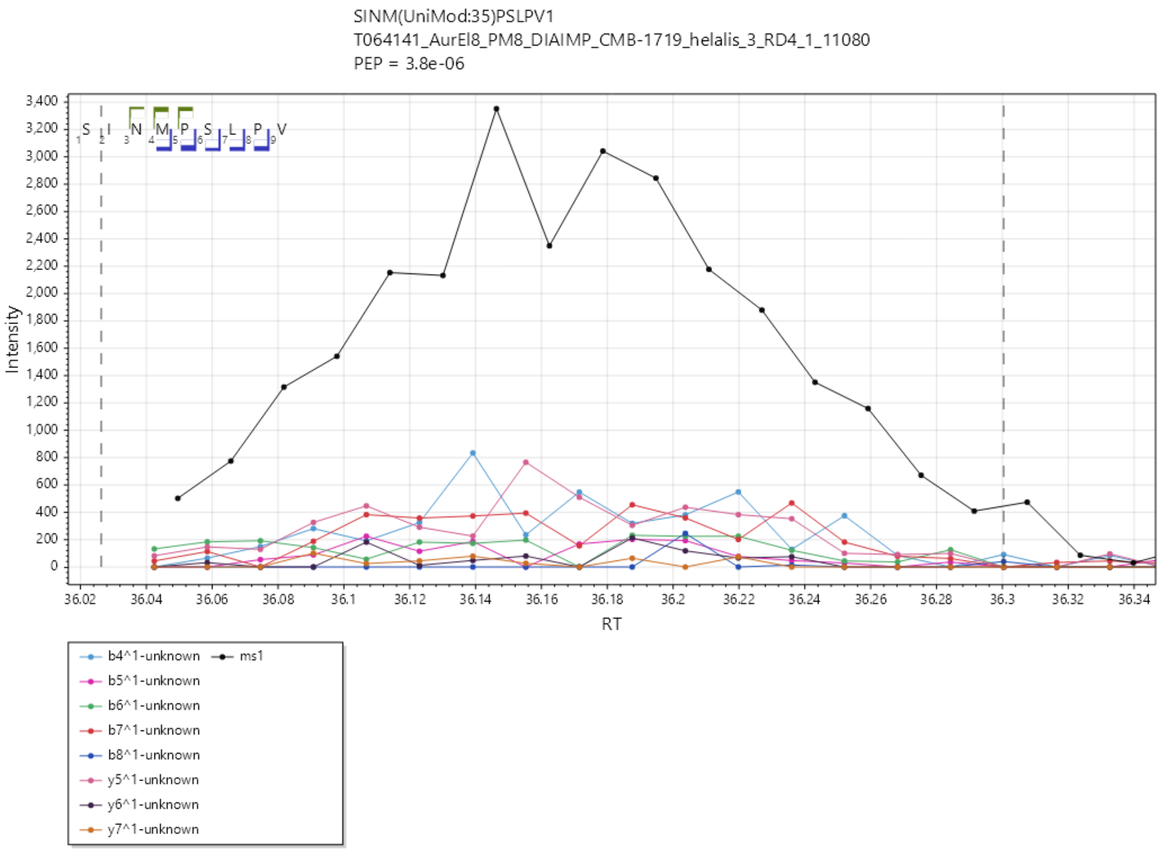

SLATIKVIGV/1+, UniProtKB A0A3Q0NG95 matching LMON_2103, ftsZ, Cell division protein FtsZ.

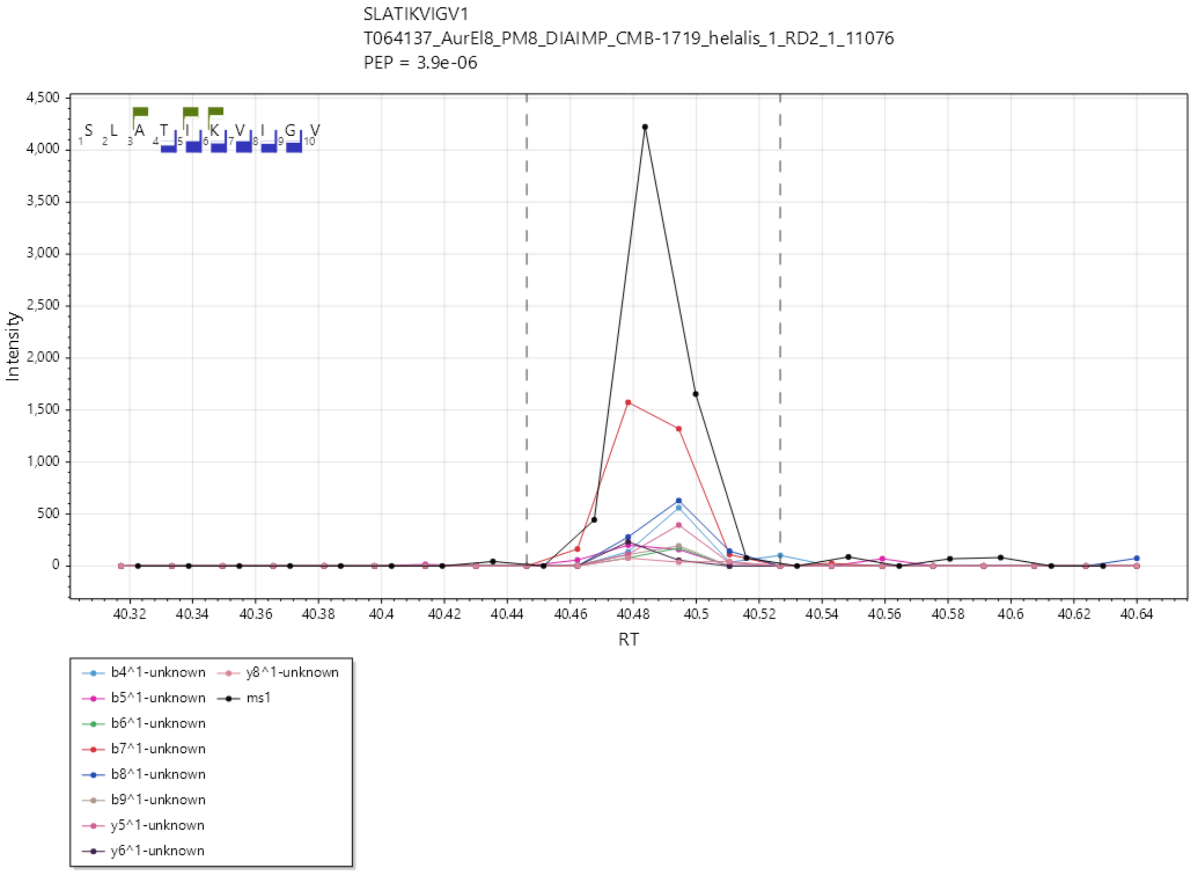

TVPGIEVIVSA/1+, UniProtKB A0A3Q0NHX2 matching LMON_2714, LMON_2714, Peptidoglycan hydrolase

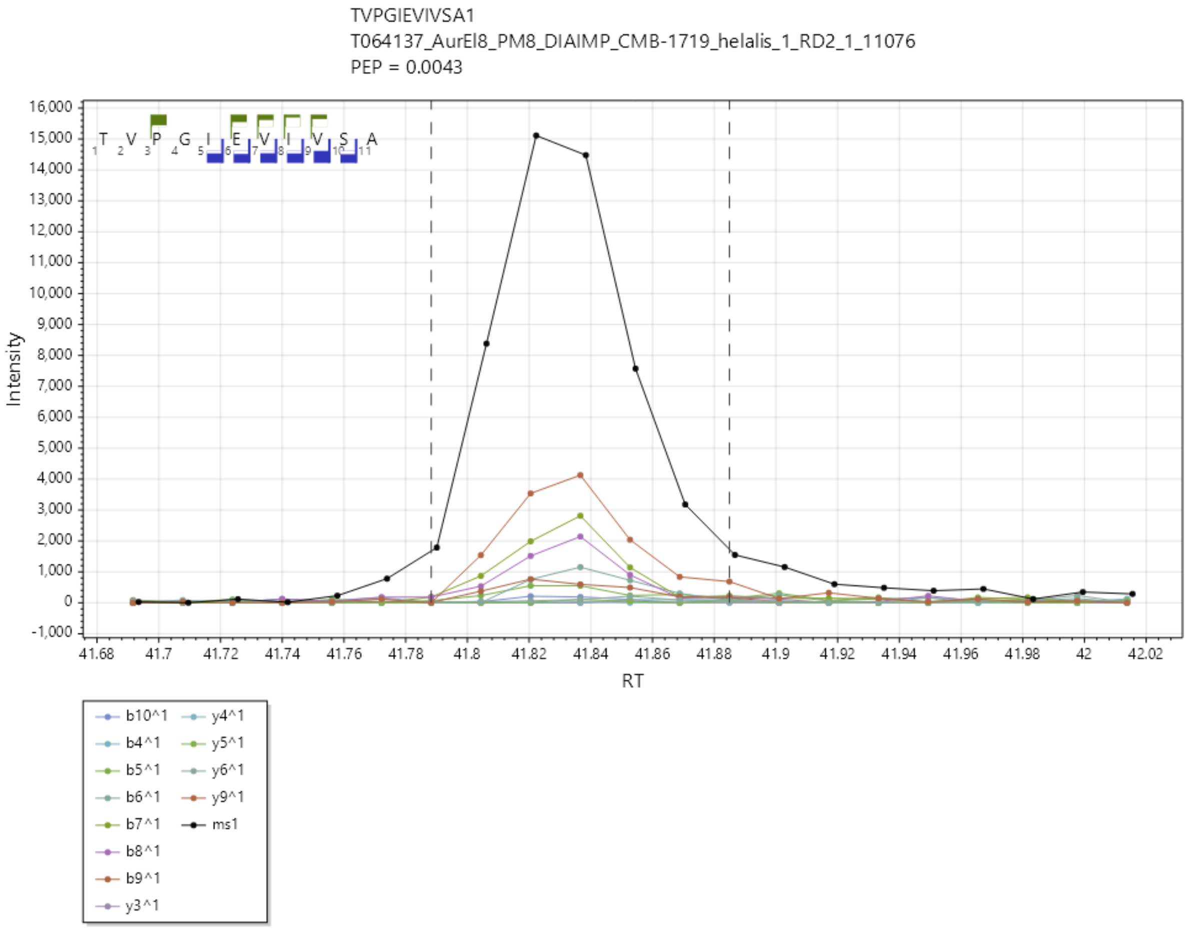

TVVAIAAGL/1+, UniProtKB A0A3Q0NHC5 matching LMON_2534, LMON_2534, Cell wall-binding protein

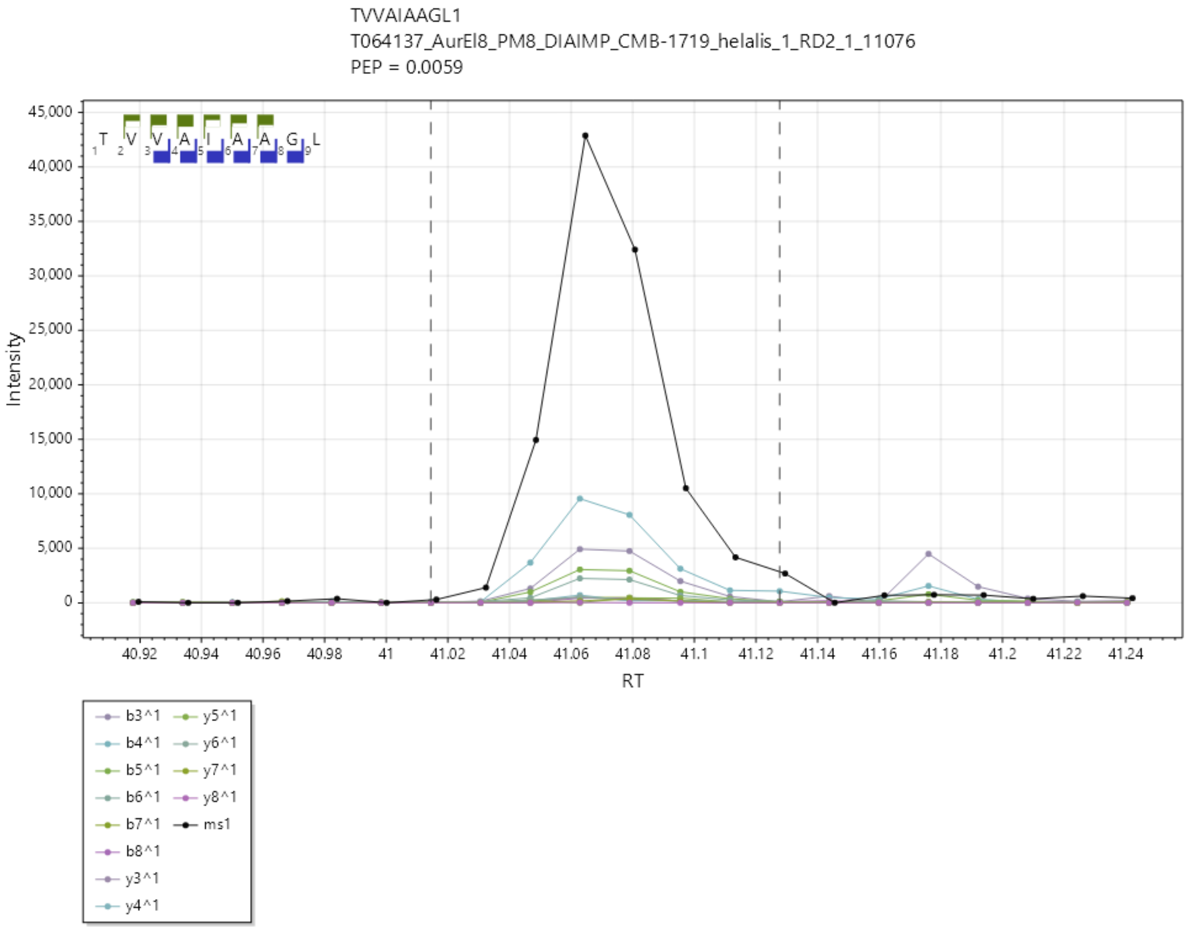

VAYGRQVYL/2+, UniProtKB A0A3Q0NAV6 matching LMON_0200, hly, Thiol-activated cytolysin.

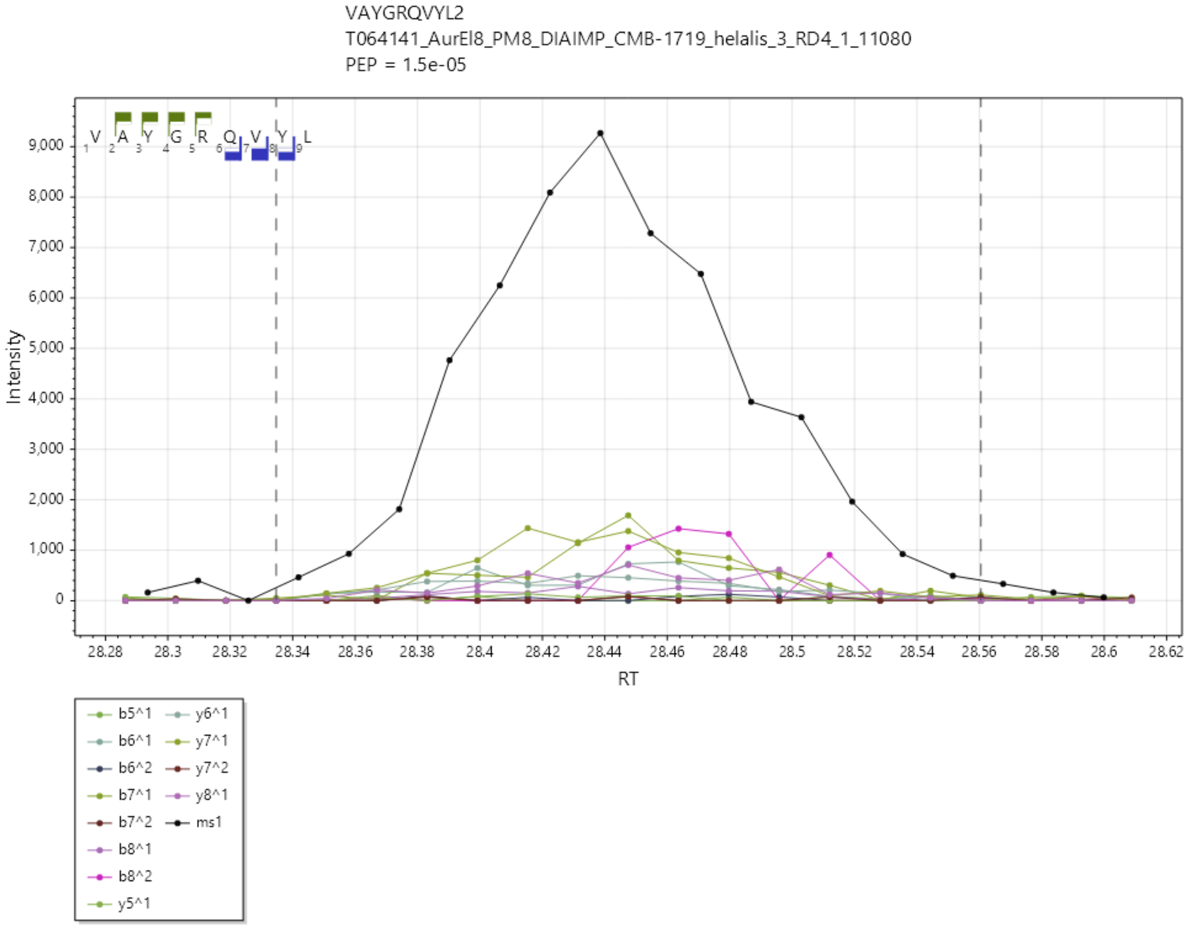

VGVPYIVVF/1+, UniProtKB A0A3Q0NHS2 matching LMON_2676, tuf, Elongation factor Tu.

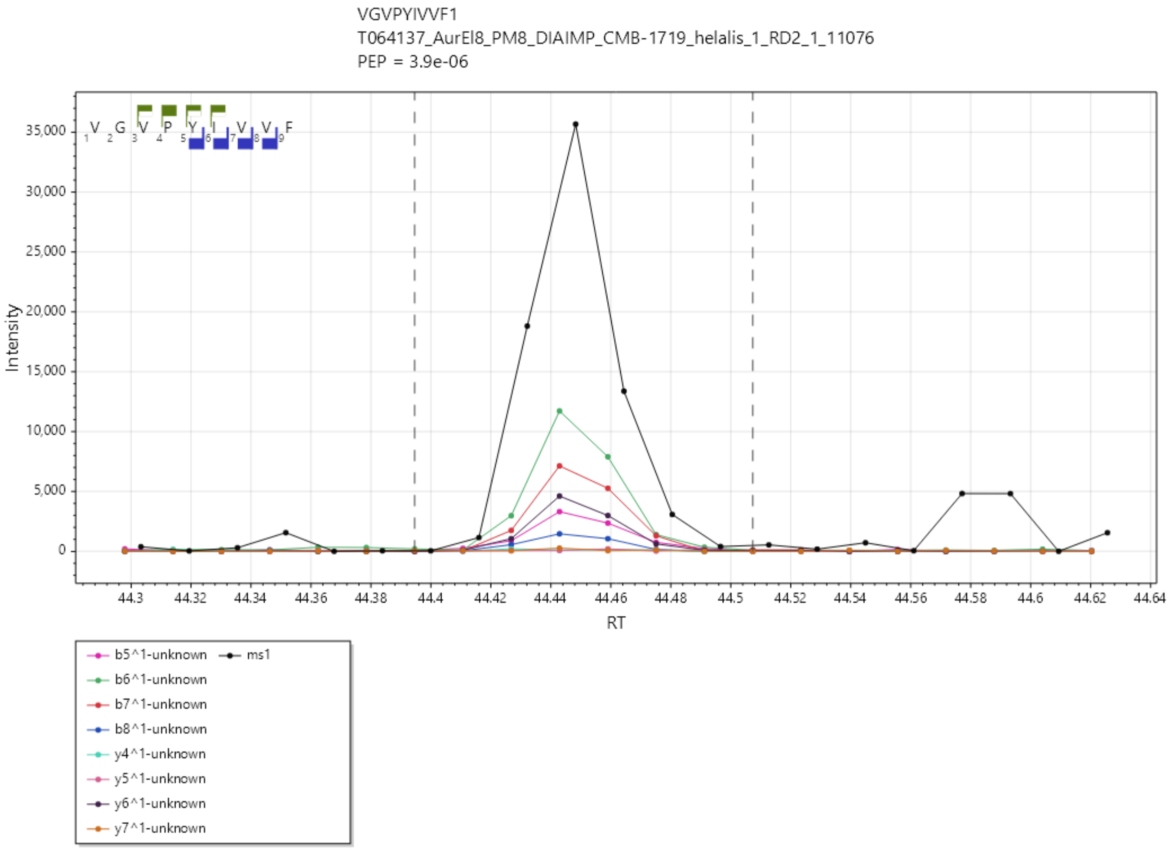

VGYSHPVEF/2+, UniProtKB A0A3Q0NJ20 matching LMON_2638, rplF, Large ribosomal subunit protein uL6.

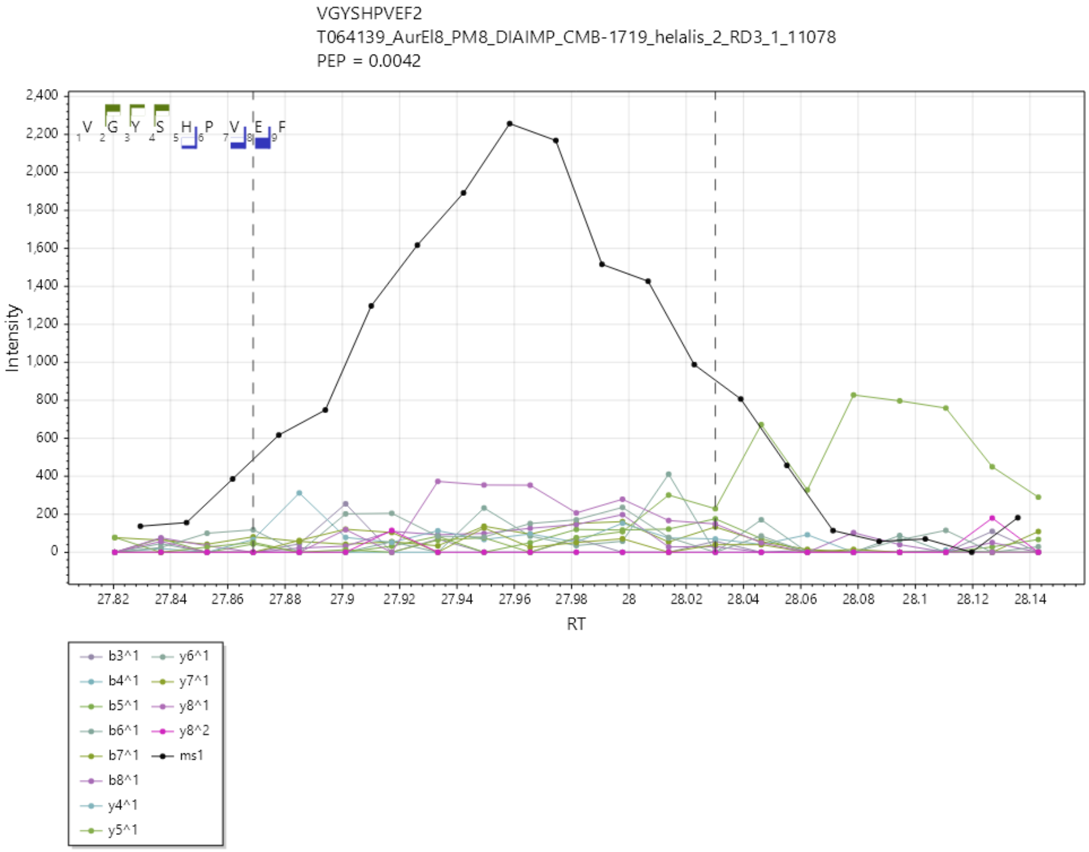

YKLGLAIHY/2+, UniProtKB A0A3Q0NAV2 matching LMON_0203 , plcB, Phospholipase C

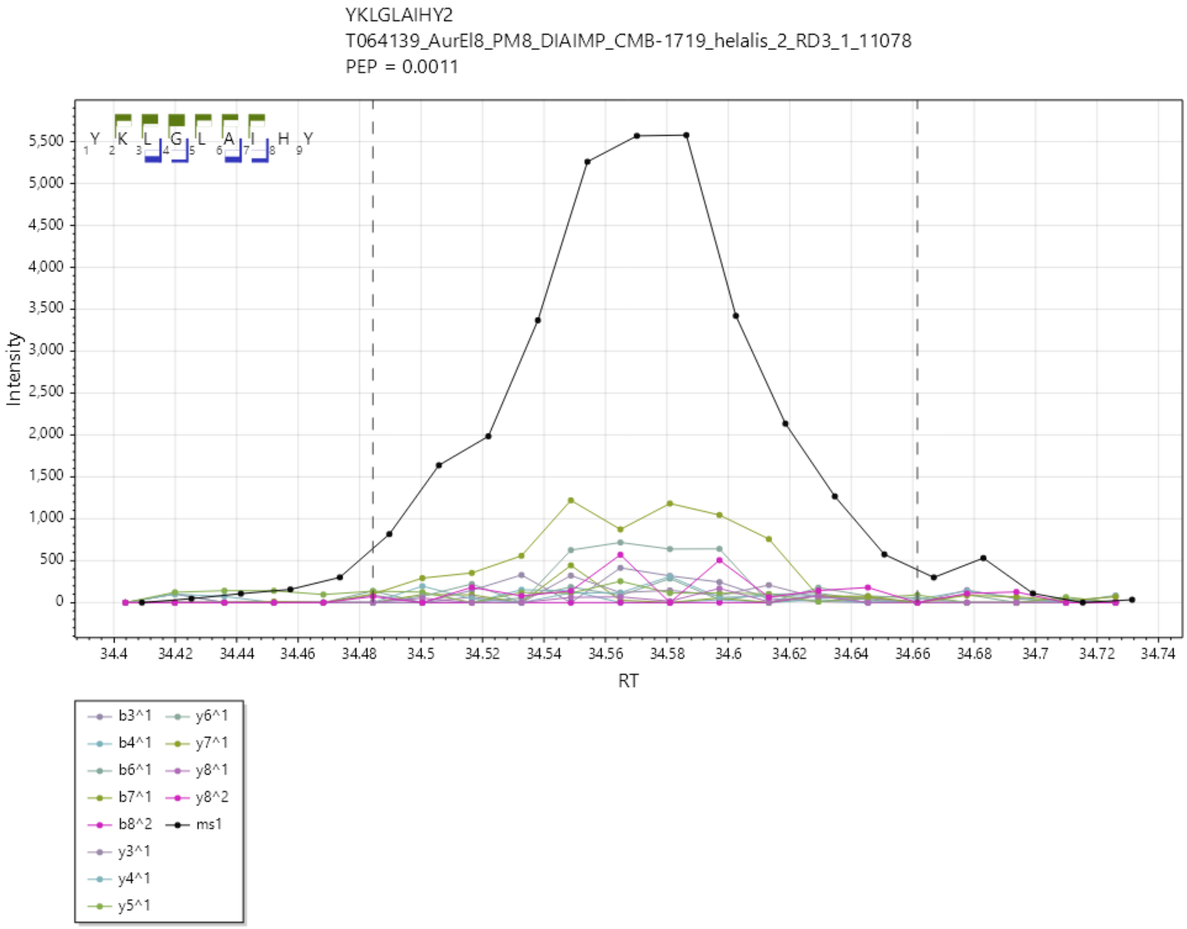

YTFEVDTRATKTQV/2+,A0A3Q0NHR6 matching LMON_2651, rplW, Large ribosomal subunit protein uL23.

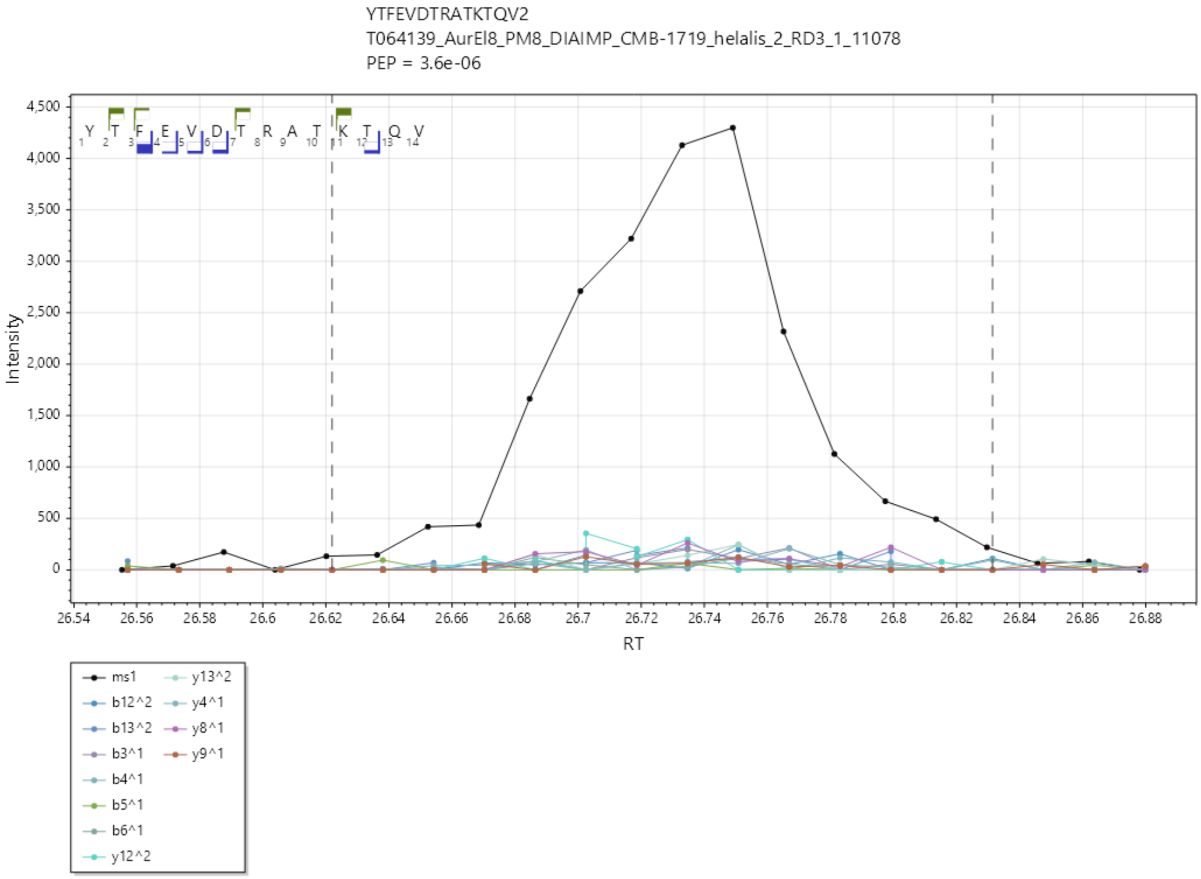
